# Multimodal subspace independent vector analysis effectively captures the latent relationships between brain structure and function

**DOI:** 10.1101/2023.09.17.558092

**Authors:** Xinhui Li, Peter Kochunov, Tulay Adali, Rogers F. Silva, Vince D. Calhoun

**Author notes:** These authors jointly supervised and equally contributed to this work (Rogers F. Silva, Vince D. Calhoun).

## Abstract

A key challenge in neuroscience is to infer the relationships between brain structure and function from high-dimensional, multimodal neuroimaging data. While conventional multivariate approaches often simplify statistical assumptions and estimate one-dimensional independent sources shared across modalities, the true relationships between latent sources are likely more complex—statistical dependence may exist both within and between modalities, and possibly span more than one dimension per modality. Here we present Multimodal Subspace Independent Vector Analysis (MSIVA), a methodology to capture both joint and unique vector sources from multiple data modalities by defining both cross-modal and unimodal subspaces with variable dimensions. In particular, MSIVA enables flexible estimation of varying-size independent subspaces within modalities and their one-to-one linkage to corresponding subspaces across modalities. A key advantage is its ability to capture subject-level variability at the voxel level within independent subspaces, in contrast to traditional methods that share the same independent components across subjects. We evaluated three initialization workflows with five candidate subspace structures in multiple synthetic datasets and two large multimodal neuroimaging datasets, both including structural MRI (sMRI) and functional MRI (fMRI). After confirming that MSIVA successfully recovered the ground-truth subspace structures in the synthetic data, we then applied MSIVA to identify the latent subspace structure in the neuroimaging data. Subsequent subspace-specific canonical correlation analysis, brain-phenotype prediction, and voxelwise brain-age delta analysis suggest that the estimated sources from MSIVA with the optimal subspace structure are strongly associated with multiple phenotype variables, including age, sex, schizophrenia, lifestyle factors, and cognitive functions. Further, we identified modality- and group-specific brain regions related to multiple phenotype measures such as age (for example, cerebellum, precentral gyrus, and cingulate gyrus in sMRI; occipital lobe and superior frontal gyrus in fMRI), sex (for example, cerebellum in sMRI, frontal lobe in fMRI, and precuneus in both sMRI and fMRI), schizophrenia (for example, cerebellar, frontal, and insular cortices in sMRI; occipital pole, lingual gyrus, and precuneus in fMRI), shedding light on phenotypic and neuropsychiatric biomarkers of linked brain structure and function.

## 1 Introduction

Neuroimaging techniques such as magnetic resonance imaging (MRI) have been developed to understand the structural and functional properties of the brain, as well as their relationships to behavior. However, it is challenging to directly associate behavior measures with raw MRI data, which typically includes tens of thousands of voxels and subjects. Although the data in its original space appears complex, its intrinsic dimensionality can be significantly lower. Recent studies have found that neural representations in low-dimensional subspaces form the basis that supports motor functions such as reaching (Churchland et al., 2012; Pandarinath et al., 2018) and timing (Remington et al., 2018; Wang et al., 2018), and cognitive functions such as perception (Bao et al., 2020; Chang & Tsao, 2017; Semedo et al., 2019; She et al., 2024), generalization (Bernardi et al., 2020; Boyle et al., 2024; Courellis et al., 2024), and decision-making (Hajnal et al., 2024; Johnston et al., 2024). Hence, it is important to develop latent variable models to learn low-dimensional representations and structures from high-dimensional data. In addition, each neuroimaging modality has its own strengths and weaknesses, and only captures limited information about the brain. For example, structural MRI (sMRI) provides high-resolution anatomical structure of the brain but does not capture temporal dynamics, while functional MRI (fMRI) measures blood-oxygenation-level-dependent (BOLD) signals over time at the cost of lower spatial resolution. Joint analysis of sMRI and fMRI can offer rich spatio-temporal information in the brain that is not captured by a single modality. With the increasing availability of multimodal neuroimaging datasets, it is necessary to develop multivariate approaches to effectively capture interpretable and multifaceted information about the brain and its disorders from multiple imaging modalities (Calhoun & Sui, 2016; Lahat et al., 2015; Sui et al., 2012; Zhang et al., 2020).

A variety of data-driven multivariate approaches have been developed to jointly analyze multiple neuroimaging datasets or data modalities, including joint independent component analysis (jICA) (Calhoun & Adali, 2008; Calhoun, Adali, Giuliani, et al., 2006; Calhoun, Adali, Pearlson, & Kiehl, 2006; Franco et al., 2008), linked ICA (Groves et al., 2011), multimodal canonical correlation analysis (mCCA) (Correa et al., 2008, 2010; Mohammadi-Nejad et al., 2017), jICA+mCCA (Sui et al., 2011, 2013), and independent vector analysis (IVA) (Adali et al., 2015a, 2015b). Notably, a unified framework Multidataset Independent Subspace Analysis (MISA) (Silva et al., 2020) has recently been introduced, encompassing multiple latent variable models, such as ICA (Comon, 1994), IVA (Adali et al., 2014; Kim et al., 2006), and independent subspace analysis (ISA) (Cardoso, 1998; Hyvärinen & Hoyer, 2000; Szabó et al., 2012). MISA can be applied to identify latent sources from multiple neuroimaging modalities, including sMRI and fMRI (Silva et al., 2020). More recently, a multimodal IVA (MMIVA) fusion method built upon MISA has been proposed to identify linked biomarkers related to age, sex, cognition, and psychosis in two large multimodal neuroimaging datasets (Silva et al., 2024). Many existing approaches, including MMIVA, assume that sources are one-dimensional and independent within each modality, that is, the subspace structure is an identity matrix. However, brain networks are hierarchically organized across multiple spatial and temporal scales (Betzel & Bassett, 2017; Felleman & Van Essen, 1991; Friston, 2008; Kiebel et al., 2008; Zeki & Shipp, 1988). This hierarchical organization implies that the underlying relationships between true latent sources are likely more complex—statistical dependence may exist both within and across modalities, and span more than one dimension per modality. Sources from the same modality may be linked, potentially grouped by their anatomical or functional properties. For instance, spatial dependencies between corresponding sources have been observed in task-based and resting-state fMRI studies (Ma et al., 2010, 2011; McKeown et al., 1998), which MMIVA would not optimally capture.

Aiming to better detect the statistical relationships from multimodal data, we present a novel methodology, Multimodal Subspace Independent Vector Analysis (MSIVA)^1^, that captures linkage of vector sources by defining cross-modal and unimodal subspaces with variable dimensions. MSIVA is built upon MMIVA by defining a block diagonal matrix as the subspace structure, instead of the identity matrix used in MMIVA. We default the initialization of MSIVA to the weight matrices learned by multimodal group principal component analysis (MGPCA) and unimodal ICAs combined. MSIVA can simultaneously estimate two types of latent sources—those linked across all modalities (along with their underlying relationships) and those unique to a specific modality. Moreover, by leveraging higher-dimensional subspaces, MSIVA sources show greater representation power, which supports downstream analyses at both individual and voxel levels.

To comprehensively assess the effectiveness of MSIVA, we compared three different initialization strategies across five candidate subspace structures. We first simulated multiple synthetic datasets to evaluate whether MSIVA can successfully reconstruct the ground-truth subspace structures. Next, we applied MSIVA to two large multimodal neuroimaging datasets, the UK Biobank dataset (Miller et al., 2016) and a schizophrenia (SZ) patient dataset combined from several studies (Aine et al., 2017; Keator et al., 2016; Tamminga et al., 2014). Using canonical correlation analysis (CCA) (Hotelling, 1992), we conducted a follow-up assessment of each MSIVA cross-modal subspace separately and identified projections within the optimal subspace structure yielding the post-CCA linked sources. We then performed age regression, sex classification, and diagnosis classification to investigate the associations between these linked sources and phenotype measures. Furthermore, we proposed a voxelwise brain-age delta analysis using reconstructed data from MSIVA.

Simulation results showed that MSIVA successfully recovered the ground-truth subspace structures including high-dimensional subspaces. In two independent neuroimaging datasets, the optimal latent structure identified by MSIVA consisted of five two-dimensional cross-modal subspaces, instead of the one-dimensional independent subspaces used in MMIVA. Brain-phenotype modeling revealed the post-CCA MSIVA sources were related to age, sex, and SZ-related effects, while the brain-age gap was explained by several phenotype measures, including lifestyle factors and cognitive test scores. Overall, our findings suggest that MSIVA can effectively reveal linked multidimensional latent sources related to phenotype variables from multimodal neuroimaging data, thereby uncovering linked phenotypic and neuropsychiatric biomarkers of brain structure and function.

## 2 Methods

### 2.1 Multimodal subspace independent vector analysis

We consider the following problem that each observed data modality is a linear mixture of latent sources:

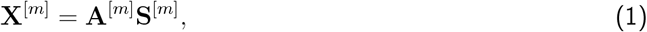

where **X**^[*m*]^ ∈ ℝ^*V* ×*N*^ is the observed data, **A**^[*m*]^ ∈ ℝ^*V* ×*C*^ is a linear mixing matrix, **S**^[*m*]^ ∈ ℝ^*C*×*N*^ is the latent source matrix, *m* is the modality index, *V* is the input feature dimensionality, *C* is the number of latent sources (*C < V*), and *N* is the number of samples. Sources across *M* modalities are either statistically dependent or independent, according to the subspace structure *S* defined using available *a priori* information. We aim to recover the latent sources **Ŝ**^[*m*]^ ∈ ℝ^*C*×*N*^ by estimating a linear unmixing matrix **W**^[*m*]^ ∈ ℝ^*C*×*V*^ :

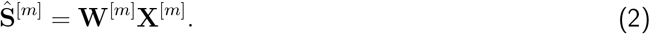

We refer to our proposed approach as Multimodal Subspace Independent Vector Analysis (MSIVA) because it extends MMIVA, linking equal size, *higher-dimensional* subspaces across modalities. We consider five candidate subspace structure priors that define different types of multimodal relationships (Figure 1) and three initialization workflows that capture different amounts of joint cross-modal information (Figure 2). Given a candidate subspace structure, MSIVA consists of iterative combinatorial optimization of the source estimates (cross-modal subspace alignment) and numerical optimization of the MISA loss (Equation 6). This process is repeated for each of the five candidate subspace structures, followed by a best-fit determination based on the final quantitative metrics of all candidates.

**Figure 1:**
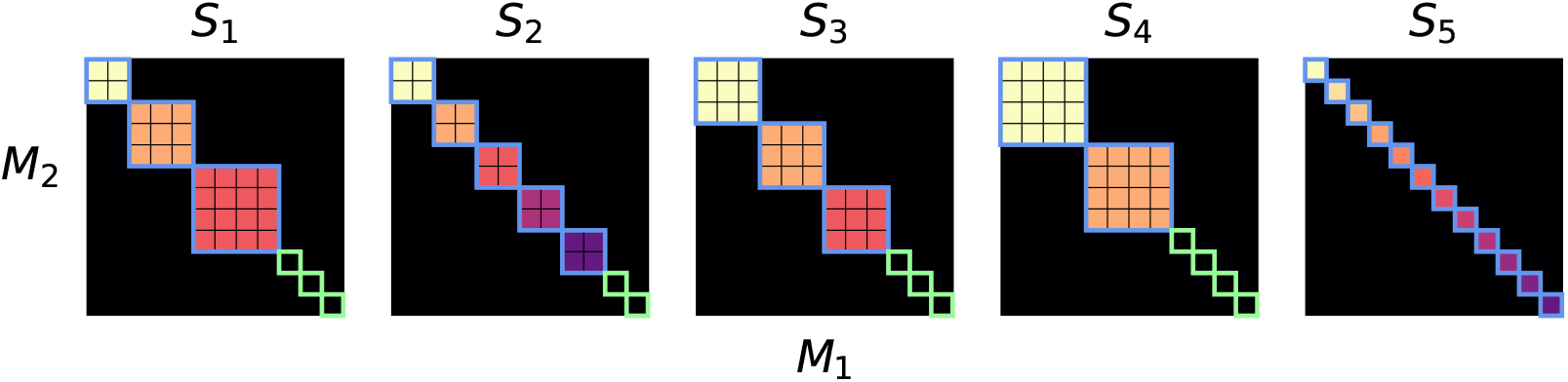
Five plausible candidate subspace structure priors (*S*_1_ − *S*_5_) for two modalities (*M*_1_, *M*_2_). Each panel depicts the idealized association between sources from two modalities (*M*_1_, *M*_2_), across five different plausible scenarios (*S*_1_ −*S*_5_). The size of each block represents the number of sources within a subspace (the subspace size). The colorful subspaces highlighted in blue are linked between modalities, whereas the black subspaces highlighted in green (1 × 1 blocks in *S*_1_ − *S*_4_) are specific to each modality (no cross-modal correlation). For each modality, sources within the same subspace are statistically dependent while sources in different subspaces are statistically independent.

**Figure 2:**
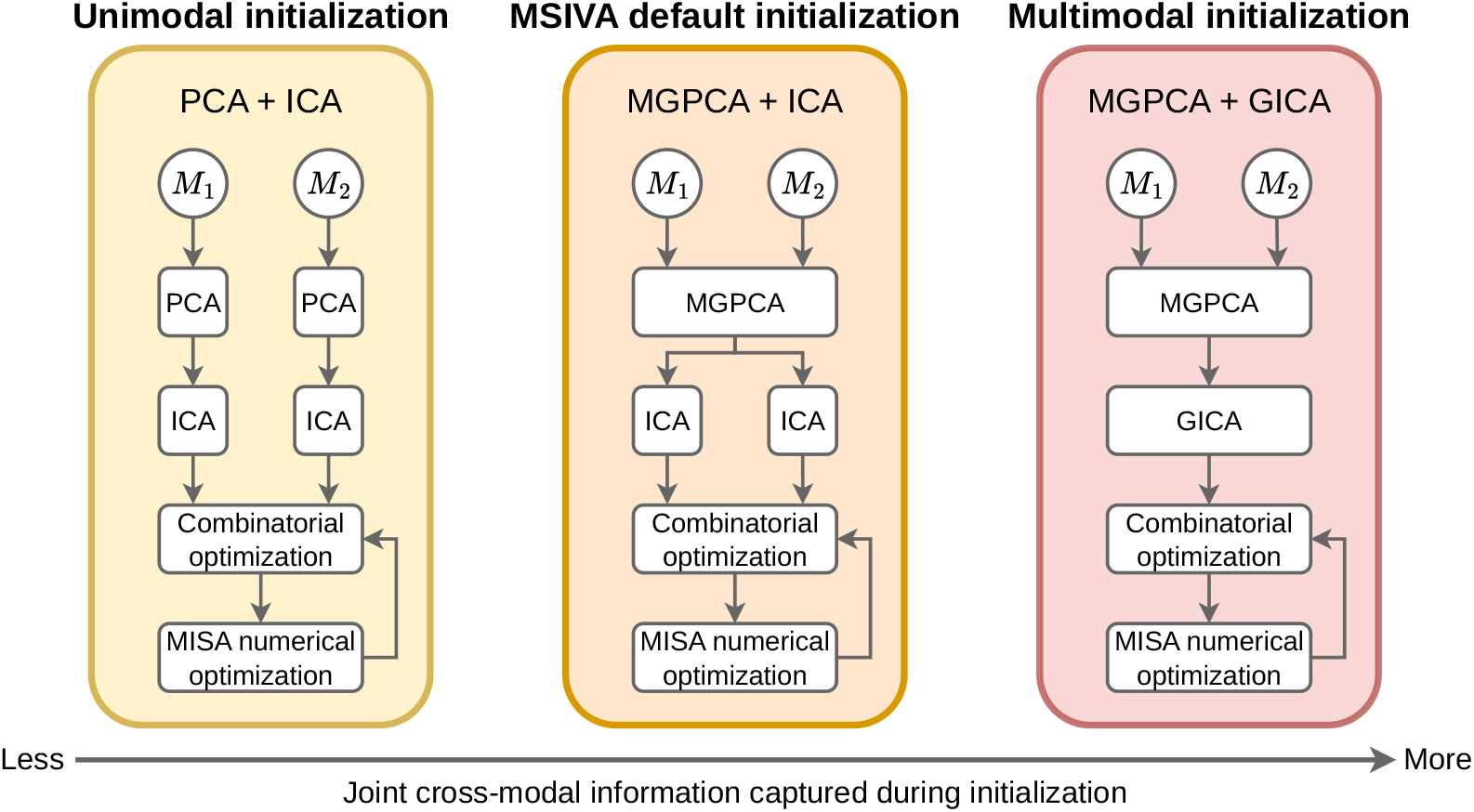
Overview of three proposed initialization workflows. From left to right, the unimodal, MSIVA default, and multimodal initialization approaches are: separate PCAs followed by separate ICAs (PCA + ICA); multimodal group PCA with separate ICAs per modality (MGPCA + ICA); multimodal group PCA with group ICA (MGPCA + GICA). MGPCA + ICA was used as the MSIVA default initialization workflow. After initialization, the greedy combinatorial optimization and the numerical optimization with the MISA loss were performed for sufficient iterations until the loss value converged.

#### 2.1.1 Designing subspace structure priors

Higher-dimensional subspaces are more likely to explain complex variability in the data, such as statistical dependence. In designing subspace structures, we consider two main objectives. First, unlike MMIVA, which assumes all sources are statistically independent, our approach aims to identify groups of linked (*not* independent) sources within each modality, while assuming independence between groups. These groups of linked sources are referred to as *subspaces*. Second, we aim to detect statistical dependence between subspaces across different modalities, which involves solving an expensive combinatorial optimization problem. To simplify this process, we restrict the search space by assuming that cross-modal dependence occurs only between higher-dimensional subspaces (two-dimensional or greater) of the same size across modalities. Additionally, we treat all modality-specific subspaces as one-dimensional (1*D*), meaning each represents a single unlinked unimodal source.

Building on the MISA framework (Silva et al., 2020), we require user-defined candidate subspace structure priors that specify the expected linkage patterns. The goal of MSIVA is to determine which one of the candidate subspace structures best fits the observed data. We consider two questions when designing the subspace structure. First, *how many sources should be included?* We recommend estimating the model order of the real data using information-theoretic criteria (Y.-O. Li et al., 2007) and selecting the number of sources that captures sufficient information on all modalities. Second, *how should the sources be grouped?* In the functional imaging literature, two-to four-dimensional clusters have been used to hierarchically group sources (Ke et al., 2019; Ma et al., 2010, 2011). Accordingly, we define subspace structures with two-to four-dimensional cross-modal subspaces. We consider subspace structures that include as many groups of uniform dimension as possible, as well as one subspace configuration that contains groups of different dimensions. Therefore, we propose five plausible subspace structures (*S*_1_−*S*_5_) in two modalities (*M*_1_, *M*_2_), all with 12 sources in each modality (Figure 1):

- *S*_1_: One two-dimensional (2*D*) cross-modal subspace, one three-dimensional (3*D*) cross-modal subspace, one four-dimensional (4*D*) cross-modal subspace, and three 1*D* unimodal subspaces.
- *S*_2_: Five 2*D* cross-modal subspaces and two 1*D* unimodal subspaces.
- *S*_3_: Three 3*D* cross-modal subspaces and three 1*D* unimodal subspaces.
- *S*_4_: Two 4*D* cross-modal subspaces and four 1*D* unimodal subspaces.
- *S*_5_: Twelve 1*D* cross-modal subspaces (no unimodal subspaces, identical to MMIVA).

#### 2.1.2 MSIVA default initialization workflow

Our previous research demonstrates that proper initialization of unmixing weights can pre-align latent subspaces, leading to faster convergence to optimal solutions (X. Li, Khosravinezhad, et al., 2023; Silva et al., 2024). Based on these findings, we evaluated three initialization workflows incorporating different amounts of shared cross-modal information.

MSIVA default initialization workflow first utilizes multimodal group principal component analysis (MG-PCA) to identify common principal components across all modalities and then applies unimodal ICA on the MGPCA-reduced data of each modality. Unlike principal component analysis (PCA) that identifies orthogonal directions of maximal variation for each modality separately, MGPCA identifies directions of maximal *common* variation across all modalities while ensuring equal variance contribution of each modality. Eigenvectors were computed based on the weighted average of the covariance matrices:

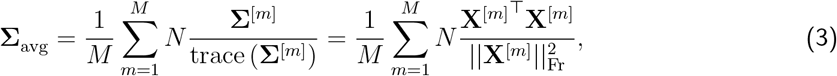

where 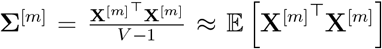, 𝔼 [·] is the expectation operator, and || · ||_Fr_ indicates the Frobenius norm. The scaling factor 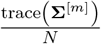 is the ratio of the variance in the modality to the number of samples. We define the whitening matrix 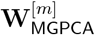 as follows:

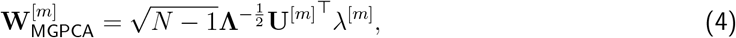

where **Λ** and **Q** are the top *C* eigenvalues and eigenvectors of **Σ**_avg_, respectively, 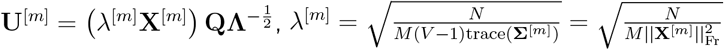.

Next, the MGPCA-reduced data from each modality 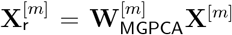 underwent a separate ICA estimation using the Infomax algorithm (Bell & Sejnowski, 1995) initialized with an identity matrix to obtain *C* independent sources per modality 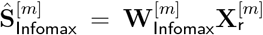. These estimates were further optimized by running MISA as a unimodal ICA model initialized with 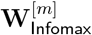, leading to the final unimodal ICA source estimates 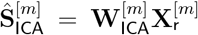. Finally, MSIVA was initialized by the combined MGPCA+ICA estimates 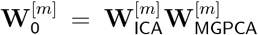 from both modalities. Subsequently, we compared MSIVA with a fully unimodal initialization workflow and a fully multimodal initialization workflow to comprehensively evaluate performance.

#### 2.1.3 Unimodal initialization workflow

The unimodal initialization workflow simply applied PCA and ICA on each modality separately. We first projected the imaging data matrix from each modality X^[*m*]^ into a reduced data matrix 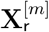 with *C* principal components and obtained the corresponding whitening matrix 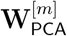 . Next, we applied ICA on each reduced data matrix 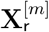 to obtain *C* independent sources and the corresponding unmixing matrix 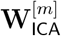 . In this workflow, the MISA initialization matrix was defined as 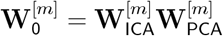.

#### 2.1.4 Multimodal initialization workflow

The multimodal initialization workflow sequentially applied MGPCA (same as in Equation 4) and group ICA (GICA) across all data modalities, resulting in the weight matrices 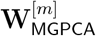 and 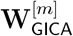. GICA performed ICA on the combined MGPCA-reduced data from all *M* modalities, that is, 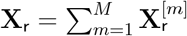. In this workflow, MISA was initialized by 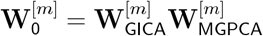.

#### 2.1.5 Alternating combinatorial and numerical optimization

All three workflows utilize MISA’s greedy combinatorial optimization and objective function to estimate latent sources. MISA uses the relative gradients and the L-BFGS algorithm (Liu & Nocedal, 1989) in a barrier-type numerical optimizer (`fmincon` from MATLAB Optimization Toolbox)^2^. The MISA loss function ℒ_MISA_(·) (Silva et al., 2020) is defi(ne)d as the Kullback-Leibler (KL) divergence between the joint distribution of all recovered sources *p* (_**Ŝ**_) and the product of all *K* subspace distributions 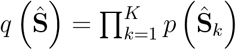, which is equivalent to mutual information among *K* subspaces. Thus, subspaces are assumed to be statistically independent of each other within each modality. Also, sources within a subspace are considered to be dependent (or linked) to one another. The distribution of each subspace is modeled as the joint multivariate Kotz distribution (Kotz, 1975; Nadarajah, 2003) of sources within that subspace as follows:

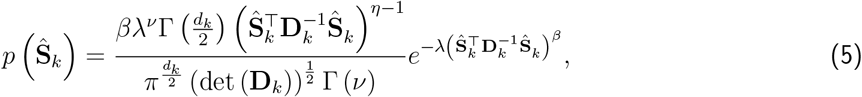

where *d*_*k*_ is the dimensionality of the *k*-th subspace (*d*_*k*_ = *C*_*k*_ · *M* if the subspace is cross-modal—*C*_*k*_ being the subspace size in each modality—and *d*_*k*_ = 1 if the subspace is unimodal). *β >* 0, *λ >* 0, and 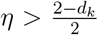 are the Kotz parameters that control the shape of the distribution, the kurtosis, and the hole size, respectively. For brevity, we define 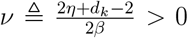 and 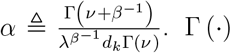 denotes the gamma function. det (·) denotes the determinant. The positive definite dispersion matrix **D**_*k*_ is given by **D**_*k*_ = *α*^−1^**Σ**_*k*_, where **Σ**_*k*_ is the source covariance matrix in the *k*-th subspace. In MSIVA, **Σ**_*k*_ is iteratively reset as the current estimate of source *correlation* matrix for subspace *k*, while the Kotz parameters are set to *β* = 0.5, *λ* = 0.8966, *η* = 1, producing a distribution slightly less peaked than a multivariate Laplace distribution with source variance 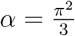, corresponding to the variance of the successful Logistic distribution used in Infomax.

We want to minimize the loss function ℒ_MISA_(·) by solving the following optimization problem:

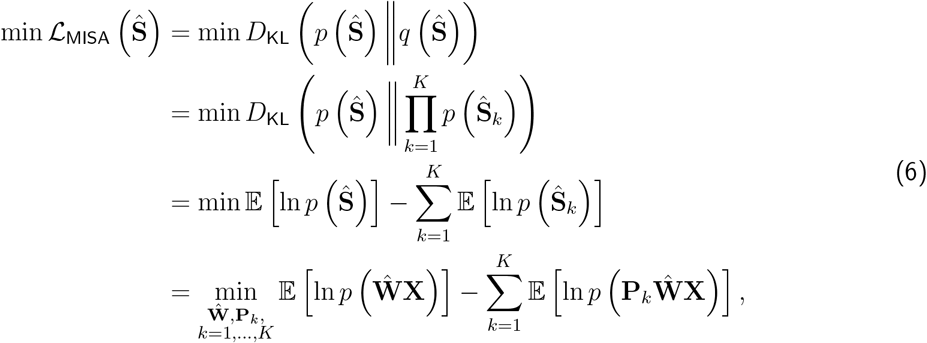

where **Ŝ**= [**Ŝ**^[1]^; … ; **Ŝ**^[*M*]]^∈ ℝ^*MC*×*N*^ represents all estimated sources from all *M* modalities^3^. **X** = [**X**^[1]^; … ; **X**^[*M*]^] ∈ ℝ^*MV* ×*N*^ represents the concatenated data from all *M* modalities. 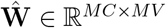 is the estimated block-diagonal unmixing matrix, such that 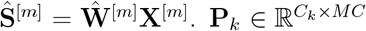 is the *k*-th subspace assignment matrix defined by the subspace structure *S* in Section 2.1.1, and *C*_*k*_ is the number of sources assigned to the *k*-th subspace.

We alternated between greedy combinatorial optimization and MISA numerical optimization until the loss value converged. Combinatorial and numerical optimizations were performed 10 times for synthetic data and 20 times for neuroimaging data. The goal of combinatorial optimization is to escape potential local minima found by the numerical optimizer. During each combinatorial optimization, the rows of the modality-specific unmixing matrices 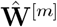 were permuted. The numerical optimizer then resumed for an additional 150 iterations, returning to combinatorial optimization afterwards. Finally, we selected the 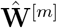 with the lowest loss value across all iterations performed.

### 2.2 Datasets

#### 2.2.1 Synthetic data

For each subspace structure *S*, we generated a synthetic dataset with two modalities **X** = **X**^[1]^; **X**^[2]^ ∈ ℝ^2*V* ×*N*^, where *V* is the size of input features (*V* = 20000) and *N* is the number of samples (*N* = 3000). *V* is a sufficiently large number for covariance estimation in PCA or MGPCA. *N* is chosen to approximate the number of samples in the UK Biobank neuroimaging dataset in this study (see Section 2.2.2). Each data modality is a linear mixture of *C* = 12 sources spanning the subspaces defined in *S*, **X**^[*m*]^ = **A**^[*m*]^**S**^[*m*]^, **A**^[*m*]^ ∈ ℝ^*V* ×*C*^, **S**^[*m*]^ ∈ ℝ^*C*×*N*^, *m* ∈ {1, 2}. Each subspace **S**_*k*_ is independently sampled from a multivariate Laplace distribution and hence, the marginal distributions correspond to the individual sources. Cross-modal sources within each linked subspace are dependent with correlation coefficients uniformly sampled from 0.65 to 0.85. Unimodal sources (1*D* subspaces in *S*_1_ − *S*_4_) also have Laplace distribution but independent from all other sources and subspaces. Thus, they do not correlate with any other sources.

#### 2.2.2 Neuroimaging data

We utilized two large multimodal neuroimaging datasets including two imaging modalities: T1-weighted structural MRI (sMRI) and resting-state functional MRI (fMRI). The first dataset is from the UK Biobank study (Miller et al., 2016). 2907 subjects from two sites (age mean ± standard deviation: 62.09 ± 7.32 years; age median: 63 years; age range: 46 − 79 years; 1452 males, 1455 females) were used for formal analysis after excluding subjects with more than 4% missing phenotype measures (Smith et al., 2015). The second dataset includes 999 patients and controls (age mean ± standard deviation: 38.61 ± 13.13 years; age median: 39 years; age range: 15 − 65 years; 625 males, 374 females; 538 controls, 337 patients diagnosed with schizophrenia, 63 patients with bipolar disorder, 11 patients with schizoaffective disorder, 28 schizoaffective bipolar-type probands, and 22 schizoaffective depression-type probands) combined across several studies, including Bipolar and Schizophrenia Network for Intermediate Phenotypes (BSNIP) (Tamminga et al., 2014), Center for Biomedical Research Excellence (COBRE) (Aine et al., 2017), Function Biomedical Informatics Research Network (FBIRN) (Keator et al., 2016), and Maryland Psychiatric Research Center (MPRC).

For each dataset, we preprocessed sMRI and fMRI to obtain the gray matter (GM) and mean-scaled amplitude of low frequency fluctuations (mALFF) feature maps, respectively. We resampled each GM or mALFF feature map to 3 × 3 × 3mm^3^ resolution and applied a group-level GM mask on the feature map, resulting in 44318 voxels. Data acquisition and preprocessing details are described in Appendix A.

Next, for each data modality in each dataset, we performed variance normalization (removed mean and divided by standard deviation) for each subject, and then removed the mean across all subjects for each voxel. Lastly, we regressed out site effects for each dataset as follows:

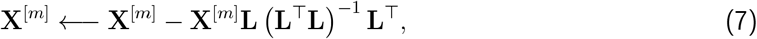

where **L** = [**1, *ℓ***], with **1** ∈ ℝ^*N*^ being a column vector of ones and ***ℓ*** being one-hot encoded site labels.

### 2.3 Quantitative evaluation metrics

#### 2.3.1 Normalized multidataset Moreau-Amari intersymbol interference

The normalized multidataset Moreau-Amari intersymbol interference (MISI) (Amari et al., 1996; Macchi & Moreau, 1995; Silva et al., 2020) was used to evaluate the residual interference between the estimated unmixing matrix 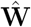 and the ground-truth mixing matrix **A**:

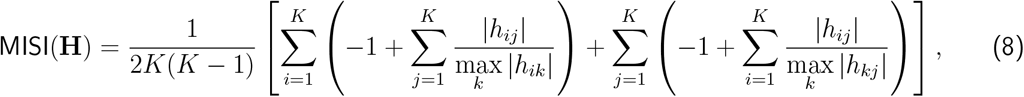

where 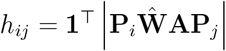 is the sum of absolute values from all elements corresponding to subspaces *i* and *j* in the interference matrix 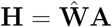.

#### 2.3.2 MISA loss

We also reported the corresponding MISA loss value defined in Equation 6. When evaluating performance on synthetic data, we prioritized the MISI metric and interference matrix **H** as they leverage the ground-truth information. We also examined whether the loss values are consistent with the MISI values.

#### 2.3.3 Mean correlation coefficient and minimum distance

We used two metrics to summarize cross-modal source correlations: the mean correlation coefficient (MCC) and the minimum distance (MD). Both metrics were computed in a two-stage manner. First, we constructed an aggregated correlation matrix across cross-modal subspaces. Then, we computed the MCC or MD on the aggregated matrix. This approach ensures balanced contributions from subspaces with varying dimensionalities. We use the term *multimodal* MCC/MD to refer to correlations computed between recovered sources and ground-truth sources within each modality, after min–max subspace aggregation. We use *cross-modal* MCC/MD to denote correlations computed between recovered sources across two different modalities, without reference to the ground truth.

The proposed metrics included three distinct correlation matrices: (1) absolute correlations between recovered sources and ground-truth sources of the first modality **R**^[1]^ ∈ ℝ^*C*×*C*^ ; (2) absolute correlations between recovered sources and ground-truth sources of the second modality **R**^[2]^ ∈ ℝ^*C*×*C*^ ; (3) absolute cross-modal correlations between recovered sources across the two modalities **R**^cm^ ∈ ℝ^*C*×*C*^ . The correlations of the first modality **R**^[1]^ were sorted into a block-diagonal structure consistent with the predefined subspace structure, and the same permutation was then applied to the correlations of the second modality **R**^[2]^. The cross-modal correlations **R**^cm^ did not involve the ground-truth, and they were left unsorted to preserve their original linkage. In the following equations, *δ*(condition) = 1 when the specified condition is satisfied.

#### Multimodal mean correlation coefficient

Given a modality-specific correlation matrix **R**^[*m*]^, we first calculated the aggregated modality-specific correlation matrix 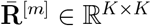 by aggregating cross-modal subspaces:

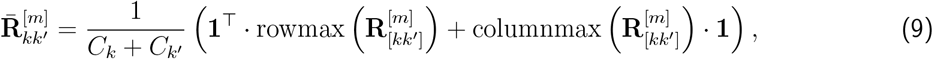

where *K* is the number of cross-modal subspaces in a given subspace structure, and the modality index 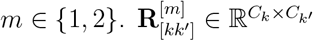 is the correlation block between the *k*-th subspace and the *k*^′^-th subspace in **R**^[*m*]^. **1** denotes a column vector of all ones. rowmax (**R**) denotes a column vector containing the maximum value of each row in the matrix **R**, and columnmax (**R**) denotes a row vector containing the maximum value of each column in **R**. *C*_*k*_ is the dimension of the *k*-th cross-modal subspace.

We then obtained the multimodal correlation matrix 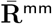 by taking the element-wise minimum for the diagonal elements and the element-wise maximum for the off-diagonal elements between the two aggregated unimodal correlation matrices (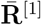 and 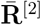):

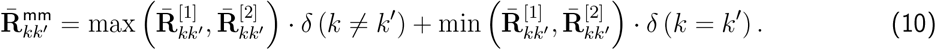

We then computed the multimodal mean correlation coefficient (MMCC) by aggregating across the diagonal of 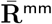:

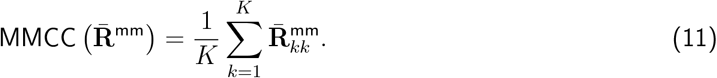

#### Cross-modal mean correlation coefficient

Similarly, we first calculated the aggregated cross-modal correlation matrix 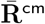:

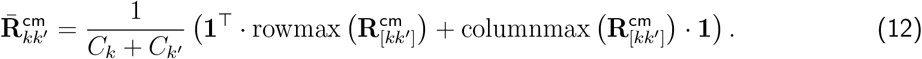

We then computed the cross-modal mean correlation coefficient (CMCC) by aggregating across the diagonal of 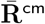:

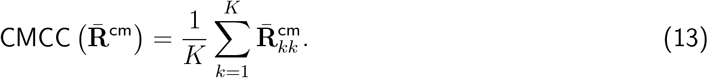

#### Multimodal and cross-modal minimum distance

We computed the multimodal minimum distance (MMD) and the cross-modal minimum distance (CMD) by aggregating *all* (diagonal and off-diagonal) elements of the matrices 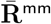 and 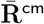, respectively. The MMD and CMD account for not only block-diagonal alignment, but also off-diagonal separation.

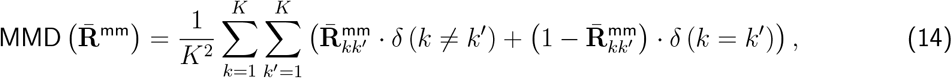

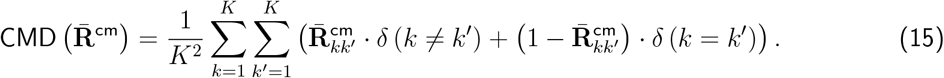

### 2.4 Experiments

#### 2.4.1 Synthetic data experiment

We first verified whether the proposed approaches including MSIVA can identify and distinguish the correct subspace structure (the one used to generate the data) from the incorrect ones in synthetic data. For each of the five subspace structures (*S*_1_ − *S*_5_) described in Section 2.1.1, we generated a synthetic dataset where the data distribution is defined by the corresponding subspace structure. Next, we conducted experiments on all combinations of five subspace structures (Figure 1) and three initialization workflows (Figure 2). Finally, we visualized the interference matrices 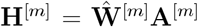^4^ to confirm if the subspace structures were recovered. In addition, we reported MISI and final MISA loss values.

#### 2.4.2 Neuroimaging data experiment

We performed experiments on each of two multimodal neuroimaging datasets separately, using each of the same five candidate subspace structures *S*_1_ − *S*_5_, and identified the optimal subspace structure as the one yielding the lowest final MISA loss value. Note that the MISI is not available because the ground-truth subspace structure is unknown in the real data. Instead, we reported the cross-modal metrics (CMCC and CMD) that measure cross-modal subspace alignment. We computed cross-modal source correlations for each subspace structure using both the *linear* Pearson correlation coefficient and the *nonlinear* randomized dependence coefficient (RDC) (Lopez-Paz et al., 2013).

To further assess the cross-modal linkage strength of the estimated subspaces *within* the optimal subspace structure, separate post-hoc CCA of each cross-modal subspace was used to recover projections with the maximum correlation between the two modalities:

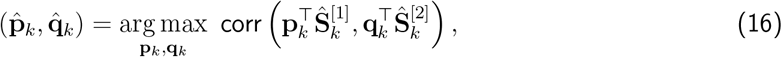

where 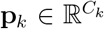 and 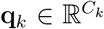are the CCA projection vectors for the *k*th cross-modal subspace, and 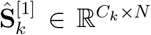 and 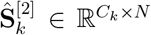 are the recovered sources in the *k*th cross-modal subspace for two modalities. After estimation, post-CCA sources in the *k*th cross-modal subspace are obtained as 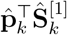 and 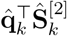. This assessment is sensible because linear transformations of individual sources within the same subspace are considered equivalently optimal^5^ (Cardoso, 1998; Szabó et al., 2012).

### 2.5 Brain-phenotype prediction

To evaluate the association between phenotype measures and cross-modal post-CCA sources, we performed age prediction and sex classification tasks for the UKB dataset, as well as age prediction and binary diagnosis classification tasks (controls versus patients with SZ) for the patient dataset. Specifically, we trained a ridge regression model to predict age and a support vector machine with a linear kernel to classify sex groups or diagnosis groups. For the UKB dataset, 2907 subjects were stratified into a training set of 2000 subjects and a holdout test set of 907 subjects. For the patient dataset, 999 subjects were stratified into a training set of 699 subjects and a holdout test set of 300 subjects in the age prediction task; 538 controls and 337 patients with SZ were grouped into a training set of 612 subjects and a test set of 263 subjects in the diagnosis classification task. We performed 10-fold cross-validation to choose the best hyperparameter (regularization parameter set: {0.1, 0.2, …, 1}) on the training set, then trained the model using all training subjects and evaluated it on the holdout test set. Age regression performance was measured by mean absolute error (MAE) between predicted age and chronological age. Sex or diagnosis classification performance was assessed via *balanced* accuracy: 0.5×(true positive rate + true negative rate).

### 2.6 Brain-age delta analysis on UK Biobank data

A key benefit of MSIVA is that the estimated multimodal sources are more expressive by leveraging higher-dimensional (≥ 2*D*) cross-modal subspaces. To demonstrate the utility of higher-dimensional subspaces, we proposed to conduct a two-stage *voxelwise* brain-age delta analysis using the UKB estimated sources from the optimal subspace structure. For each voxel in the reconstructed subspace (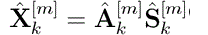^6^), we estimated an initial age delta at the first stage and corrected it for age dependence and other confound variables at the second stage (Smith et al., 2019, 2020):

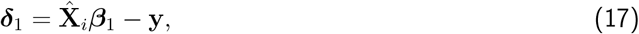

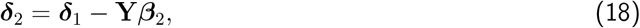

where 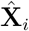 indicates the *i*-th voxel’s reconstructed patterns from each subspace. Namely, they include shared patterns from each cross-modal subspace^7^, reconstructed sMRI patterns from each cross-modal subspace, and reconstructed data from each unimodal subspace (see Appendix B for more details). **y** ∈ ℝ^*N*^ is the demeaned chronological age. **Y** ∈ ℝ^*N* ×10^ includes the confound variables: the demeaned linear, quadratic, cubic age terms, sex, the interaction between sex and each of the three age terms, the framewise displacement variable, and the spatial normalization variables from sMRI and fMRI. An advantage of the procedure described in Smith et al., 2019, 2020 is that it yields a breakdown of ***δ***_2_ per predictor in 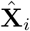. Lastly, we partialized ***δ***_2_ to remove residual associations between each predictor and the other predictors, obtaining the partialized brain-age delta, ***δ***_2*p*_.

We then correlated the voxelwise brain-age delta ***δ***_2*p*_ with 25 non-imaging phenotype variables such as lifestyle factors and cognitive test scores (see Appendix C for the full list of phenotype variables) to investigate multimodal brain-phenotype relationships. This voxelwise brain-age delta analysis allows us to visualize a voxel-level spatial map showing how each phenotype variable relates to the difference between chronological and estimated brain age.

## 3 Results

### 3.1 MSIVA identifies ground-truth subspace structures in synthetic data

We verified whether the proposed approaches, including three different initialization workflows, could identify the correct subspace structures used for data generation in synthetic datasets. As discussed in Appendix D, the MSIVA default initialization method (MGPCA+ICA) achieved a well-balanced combination of unimodal separation and cross-modal alignment in most cases. Nonetheless, after iterative combinatorial and numerical optimization, both the unimodal initialization workflow (PCA+ICA) and the MSIVA default initialization workflow led to the lowest MISI values (≤ 0.02) along the main diagonal in Figure 3, demonstrating that both approaches correctly recovered the ground truth when the correct subspace structure was provided. The multimodal initialization workflow (MGPCA+GICA), on the other hand, showed suboptimal performance with elevated MISI values along the main diagonal and was thus excluded from subsequent neuroimaging data experiments.

**Figure 3:**
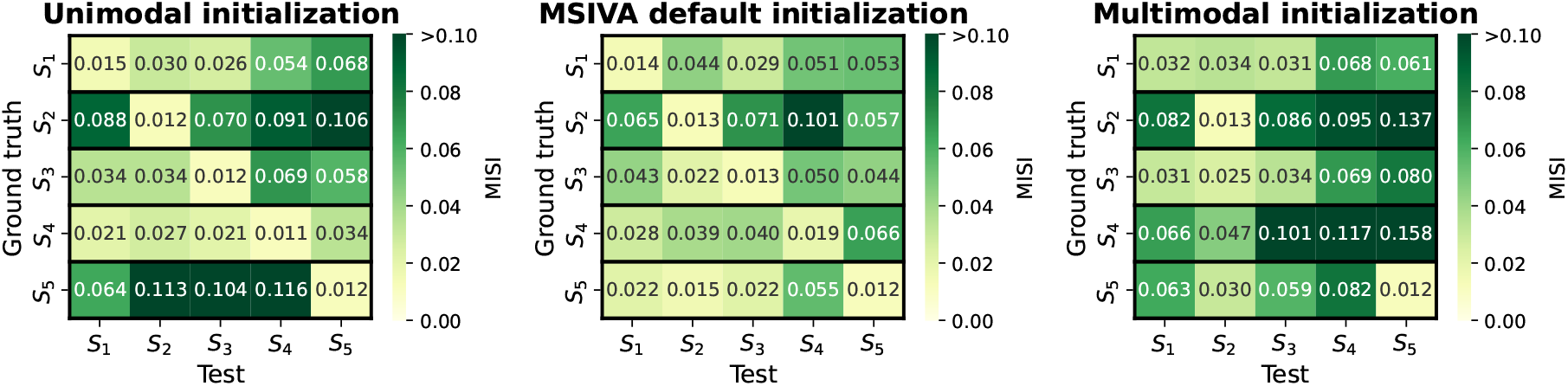
Performance on synthetic data: MISI (lower is better). Each row represents the ground-truth subspace structure used to generate the data and each column represents the test subspace structure used to fit the model. If a workflow could correctly identify all ground-truth subspace structures, the lowest MISI values would occur along the main diagonal. MISI ≤ 0.1 is considered a heuristic threshold for good source separation (Silva et al., 2014). The unimodal initialization workflow (PCA+ICA) and the MSIVA default initialization workflow (MGPCA+ICA) led to the lowest MISI values (≤ 0.02) along the main diagonal, indicating that these two initialization approaches were adequate and enabled correct identification of the ground-truth subspace structures *and* differentiation from the incorrect ones. However, the multimodal initialization workflow (MGPCA+GICA) failed to detect subspace structures *S*_1_, *S*_3_, and *S*_4_ with high MISI values in the main diagonal. Thus, the MSIVA default initialization and the unimodal initialization are considered better than the multimodal initialization.

We confirmed that loss values successfully converged within each numerical optimization run (Appendix E Figure E.1). According to Table 1, the final loss values were consistent with the MISI results. In addition, we thoroughly examined the relationship between loss values and MISI values across all numerical optimization runs. Loss values were strongly correlated with MISI values (unimodal initialization: *R*^2^ ≥ 0.95; MSIVA default initialization: *R*^2^ ≥ 0.58) when the ground-truth and test subspace structures were matched, indicating that loss serves as a valid proxy for MISI under consistent initialization (Appendix E Figure E.2). The loss values obtained with the multimodal initialization workflow failed to detect the ground-truth subspace structures containing 3*D* or 4*D* subspace(s) (*S*_1_, *S*_3_, and *S*_4_).

**Table 1:** Synthetic data: Final MISA loss values (lower is better). Each row represents the ground-truth (GT) subspace structure used to generate the data and each column represents the test subspace structure used to fit the model. The lowest loss value along the *row* is highlighted in bold, which determines the selected subspace. Approaches performing consistently well in relation to the MISI in Figure 3 would contain bold loss values *only* along the diagonal. The loss values were consistent with the MISI values. The unimodal initialization and the MSIVA default initialization correctly identified the ground-truth subspace structures. However, the multimodal initialization loss values incorrectly implied that *S*_3_, *S*_1_, and *S*_2_ were better when *S*_1_, *S*_3_, and *S*_4_ were the ground-truth subspace structures, respectively.

| Unimodal initialization | $S_1^{\text{Test}}$ | $S_2^{\text{Test}}$ | $S_3^{\text{Test}}$ | $S_4^{\text{Test}}$ | $S_5^{\text{Test}}$ |
| --- | --- | --- | --- | --- | --- |
| $S_1^{\text{GT}}$ | **42.692** | 42.879 | 42.766 | 43.054 | 43.231 |
| $S_2^{\text{GT}}$ | 42.903 | **42.301** | 42.843 | 42.823 | 42.918 |
| $S_3^{\text{GT}}$ | 42.710 | 42.864 | **42.635** | 43.073 | 43.256 |
| $S_4^{\text{GT}}$ | 43.101 | 43.238 | 43.195 | **42.975** | 43.507 |
| $S_5^{\text{GT}}$ | 43.391 | 42.997 | 43.492 | 43.767 | **42.021** |
| MSIVA default initialization | $S_1^{\text{Test}}$ | $S_2^{\text{Test}}$ | $S_3^{\text{Test}}$ | $S_4^{\text{Test}}$ | $S_5^{\text{Test}}$ |
| $S_1^{\text{GT}}$ | **42.676** | 42.880 | 42.750 | 43.002 | 43.110 |
| $S_2^{\text{GT}}$ | 42.614 | **42.288** | 42.739 | 42.876 | 42.745 |
| $S_3^{\text{GT}}$ | 42.694 | 42.848 | **42.620** | 42.868 | 43.070 |
| $S_4^{\text{GT}}$ | 33.205 | 33.275 | 33.272 | **33.115** | 33.675 |
| $S_5^{\text{GT}}$ | 43.398 | 42.946 | 43.383 | 43.969 | **42.005** |
| Multimodal initialization | $S_1^{\text{Test}}$ | $S_2^{\text{Test}}$ | $S_3^{\text{Test}}$ | $S_4^{\text{Test}}$ | $S_5^{\text{Test}}$ |
| $S_1^{\text{GT}}$ | 23.897 | 23.957 | **23.838** | 24.179 | 24.199 |
| $S_2^{\text{GT}}$ | 27.786 | **27.442** | 27.810 | 28.032 | 28.158 |
| $S_3^{\text{GT}}$ | **23.864** | 24.012 | 23.964 | 24.234 | 24.346 |
| $S_4^{\text{GT}}$ | 17.582 | **17.328** | 17.85 | 18.404 | 20.275 |
| $S_5^{\text{GT}}$ | 36.758 | 36.316 | 36.752 | 37.147 | **35.262** |

As presented in Figure 4, the recovered subspace structures from the MSIVA default initialization workflow (rows IV-V) and the unimodal initialization workflow (rows II-III) under the correct subspace structure aligned well with the proposed ground truth (row I), confirming the effectiveness of MSIVA. However, the multimodal initialization workflow (rows VI-VII) could not recover the ground-truth subspace structures for *S*_1_, *S*_3_, and *S*_4_ even when given the correct subspace structure, suggesting that cross-modal alignment optimization becomes more challenging in the presence of high-dimensional subspaces.

**Figure 4:**
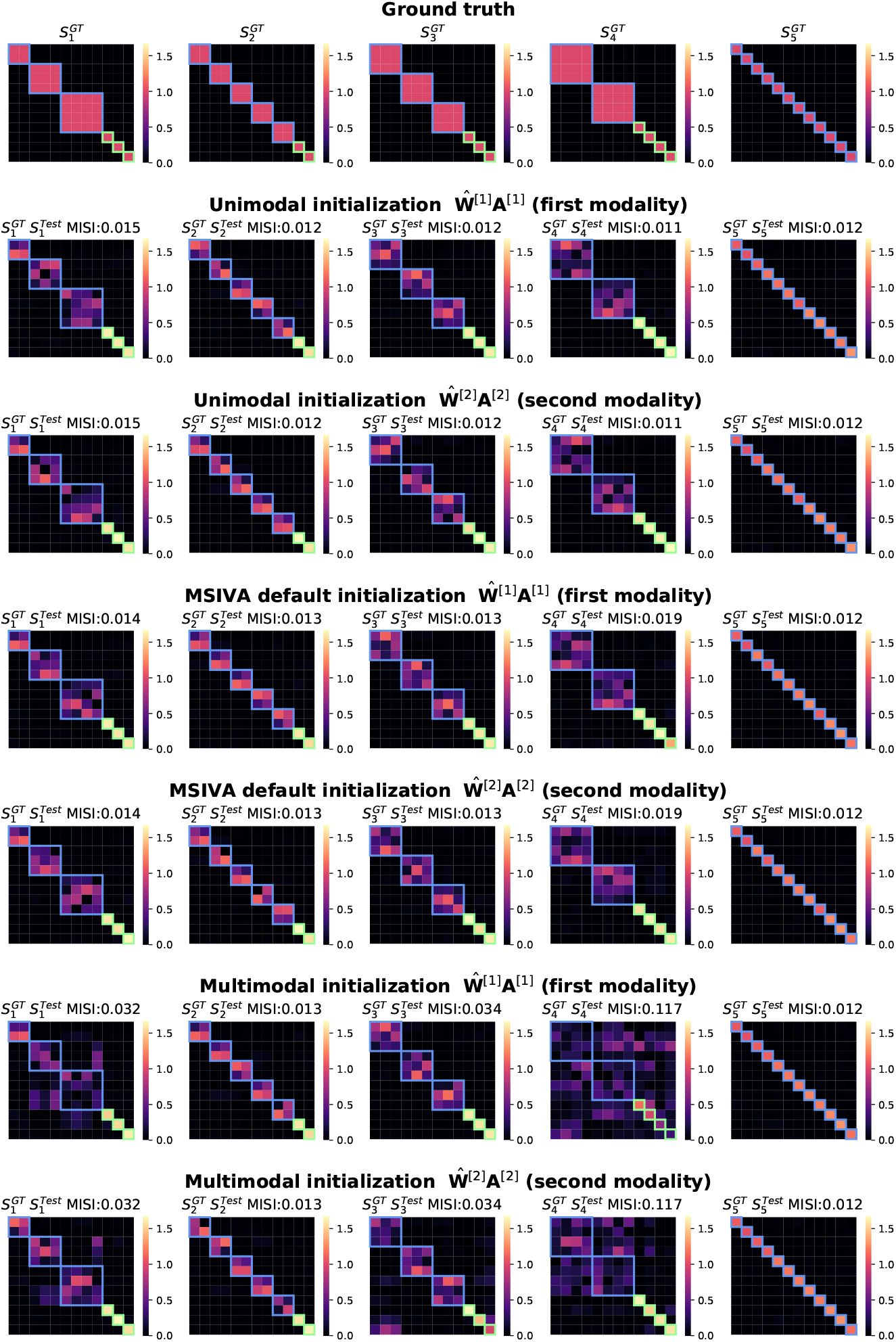
Synthetic data: Interference matrices 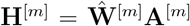 corresponding to the diagonal MISI values in Figure 3. Cross-modal subspaces are highlighted in blue while modality-specific subspaces are highlighted in green. The subspace permutation, applied for ease of interpretation, is identical for both modalities. The correct subspace structures were identified and aligned across both modalities by three workflows (rows II-VII), in accordance with the ground-truth simulation design (row I), except that the multimodal initialization failed to recover *S*_1_, *S*_3_, and *S*_4_ (rows VI-VII).

### 3.2 MSIVA detects the latent subspace structure in neuroimaging data

The model order was estimated for each neuroimaging dataset using three information-theoretic criteria (Y.-O. Li et al., 2007). Twelve sources captured the majority of information in the patient dataset and a substantial amount in the UK Biobank dataset (Appendix F). Hence, twelve sources were included in each subspace structure.

We next applied the MSIVA default initialization workflow and the unimodal initialization workflow on two large multimodal neuroimaging datasets separately—the UK Biobank (UKB) dataset and the combined patient dataset—to detect their latent subspace structures. In the UKB neuroimaging dataset, within-modal self-correlation patterns (Figure 5, rows I-II and IV-V) indicated negligible residual dependence between subspaces, as expected. Dependence within subspaces is acceptable, but dependence between subspaces is not. The MSIVA default initialization workflow recovered stronger cross-modal correlations (higher CMCCs and lower CMDs) than the unimodal initialization workflow for all predefined subspace structures (Figure 5, row VI versus row III). Results from the nonlinear dependence measure also confirmed that sources in cross-modal subspaces were linked across modalities, while sources in different subspaces within each modality were independent (Appendix G Figure G.1). Among all combinations of two initialization workflows and five candidate subspace structures, the MSIVA default initialization workflow (MGPCA+ICA) with the subspace structure *S*_2_ yielded the lowest final MISA loss value 46.775 (Table 2), suggesting that MSIVA *S*_2_ best fits the latent structure of this dataset.

**Table 2:** Neuroimaging data: Final MISA loss values (lower is better). MSIVA with the subspace structure *S*_2_ consistently yielded the lowest loss values in both multimodal neuroimaging datasets, thus it was considered as the optimal approach to capture the latent subspace structure in these two neuroimaging datasets.

| Subspace structure | $S_1$ | $S_2$ | $S_3$ | $S_4$ | $S_5$ |
| --- | --- | --- | --- | --- | --- |
| UK Biobank dataset |  |  |  |  |  |
| Unimodal initialization | 47.735 | 47.811 | 47.768 | 47.778 | 47.999 |
| MSIVA default initialization | 46.794 | **46.775** | 46.798 | 46.892 | 46.924 |
| Patient dataset |  |  |  |  |  |
| Unimodal initialization | 47.361 | 47.350 | 47.336 | 47.404 | 47.527 |
| MSIVA default initialization | 45.775 | **45.674** | 45.788 | 45.924 | 45.696 |

**Figure 5:**
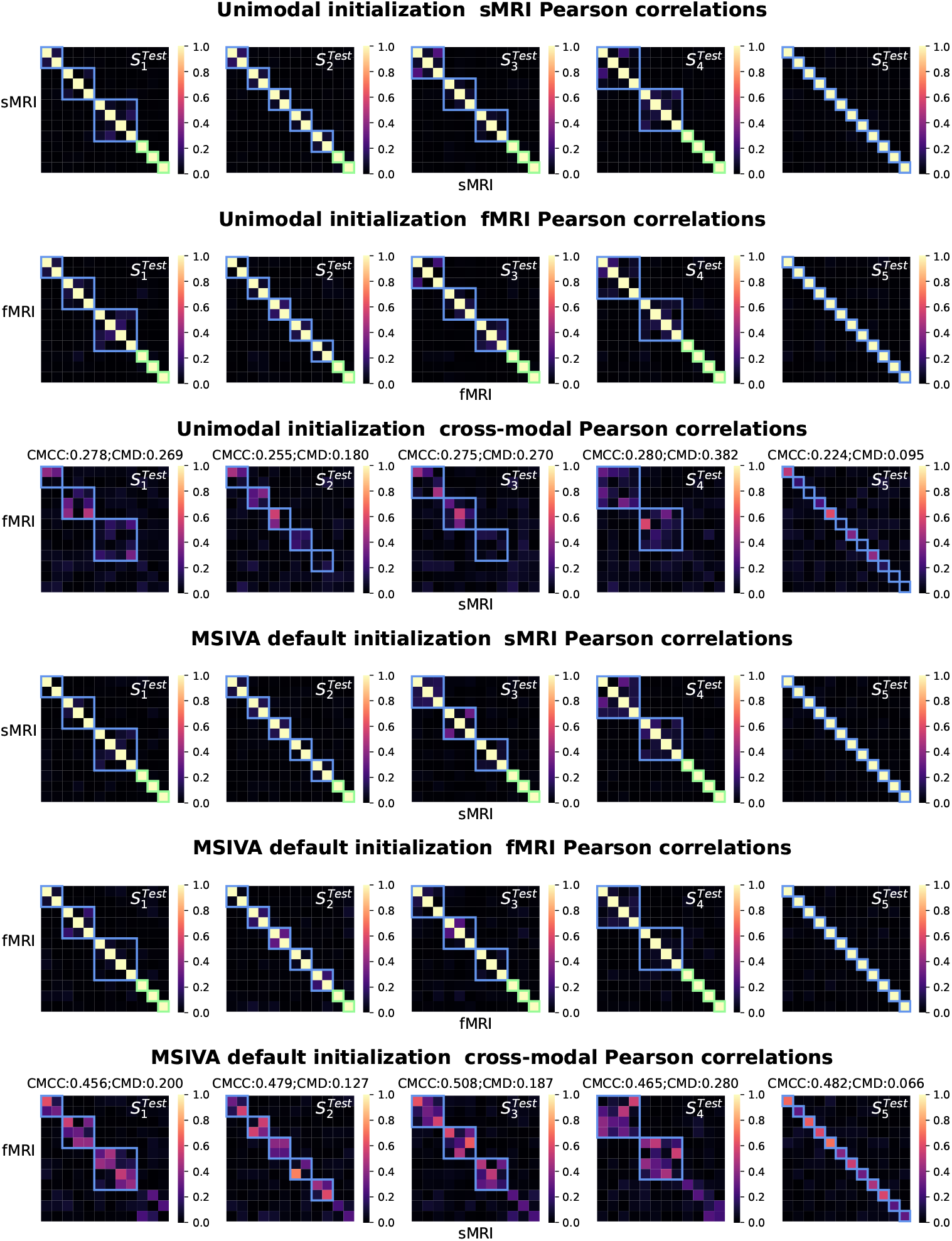
UKB neuroimaging data: Within-modal and cross-modal Pearson correlations of the recovered sources before applying post-hoc CCA. Cross-modal subspaces are highlighted in blue while unimodal subspaces are highlighted in green. Within-modal self-correlation patterns indicated negligible residual dependence between subspaces (rows I-II and IV-V). The MSIVA default initialization workflow showed stronger cross-modal correlations (higher CMCCs and lower CMDs) than the unimodal initialization workflow for all predefined subspace structures (row VI versus row III).

Similarly, in the patient dataset, the MSIVA default initialization workflow showed stronger cross-modal correlations (dependence) for all five subspace structures (Figures 6 and G.2, row VI versus row III). Same as the UKB dataset, MSIVA *S*_2_ yielded the lowest final loss value 45.674 in all cases (Table 2). These results imply that MSIVA, using the default initialization and the subspace structure *S*_2_ (including five linked 2*D* subspaces), provides a better fit to the statistical relationships in these datasets.

**Figure 6:**
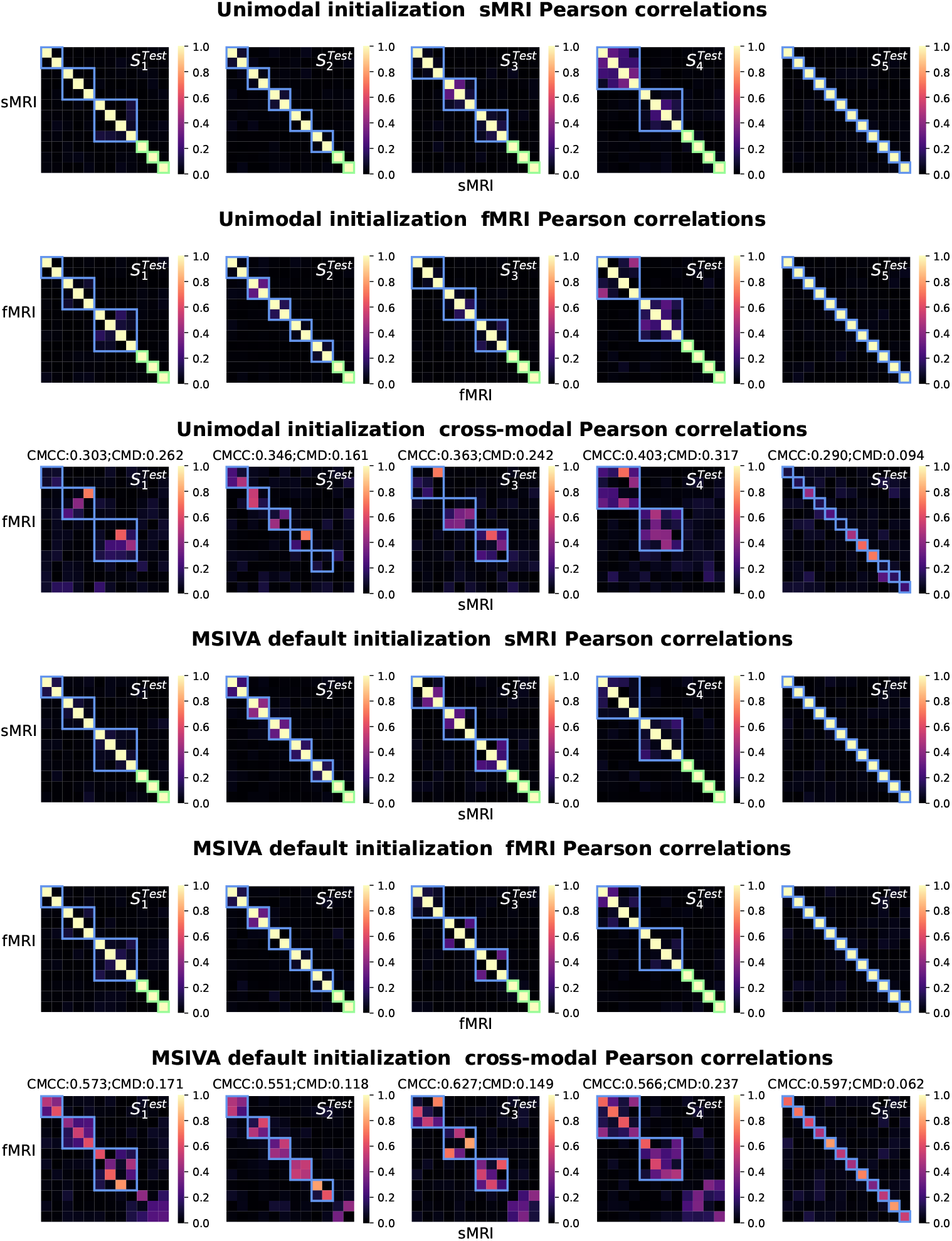
Patient neuroimaging data: Within-modal and cross-modal Pearson correlations of the recovered sources before applying post-hoc CCA. Cross-modal subspaces are highlighted in blue while unimodal subspaces are highlighted in green. Within-modal self-correlation patterns indicated negligible residual dependence between subspaces (rows I-II and IV-V). The MSIVA default initialization workflow showed stronger cross-modal correlations (higher CMCCs and lower CMDs) than the unimodal initialization workflow for all predefined subspace structures (row VI versus row III).

### 3.3 MSIVA reveals linked phenotypic and neuropsychiatric biomarkers

After identifying the neuroimaging sources, we asked whether the linked sources were biologically meaningful. To answer this question, we evaluated the brain-phenotype relationships between phenotype variables and neuroimaging sources estimated by MSIVA (with the optimal subspace structure *S*_2_ selected based on Table 2). In the UKB dataset, the cross-modal CCA projections in each linked subspace showed that subspaces 1, 3, 4, and 5 were significantly associated with age groups, particularly crossmodal source 9 in subspace 5 (Figure 7 rows I and II). Subspaces 1, 2, 3, and 4 showed significant sex differences, especially cross-modal source 7 in subspace 4 (Figure 7 rows III and IV). Furthermore, we used the post-CCA sources from each linked subspace to predict age and sex. The age regression and sex classification performance also confirmed that subspace 5 was strongly associated with age while subspace 4 was strongly associated with sex. More specifically, the age prediction MAE in subspace 5 was the lowest (5.378 years), and the sex classification balanced accuracy was the highest in subspace 4 (79.933%). In the patient dataset, the cross-modal CCA projections in each linked subspace revealed significant age effects in the cross-modal sources 1, 3, 8, 9, and 10, as well as SZ-related effects in both sources 9 and 10 from subspace 5 (Figure 8). These associations were verified by the age regression and diagnosis classification results. The sex effect in the patient dataset was not significant (Appendix H).

**Figure 7:**
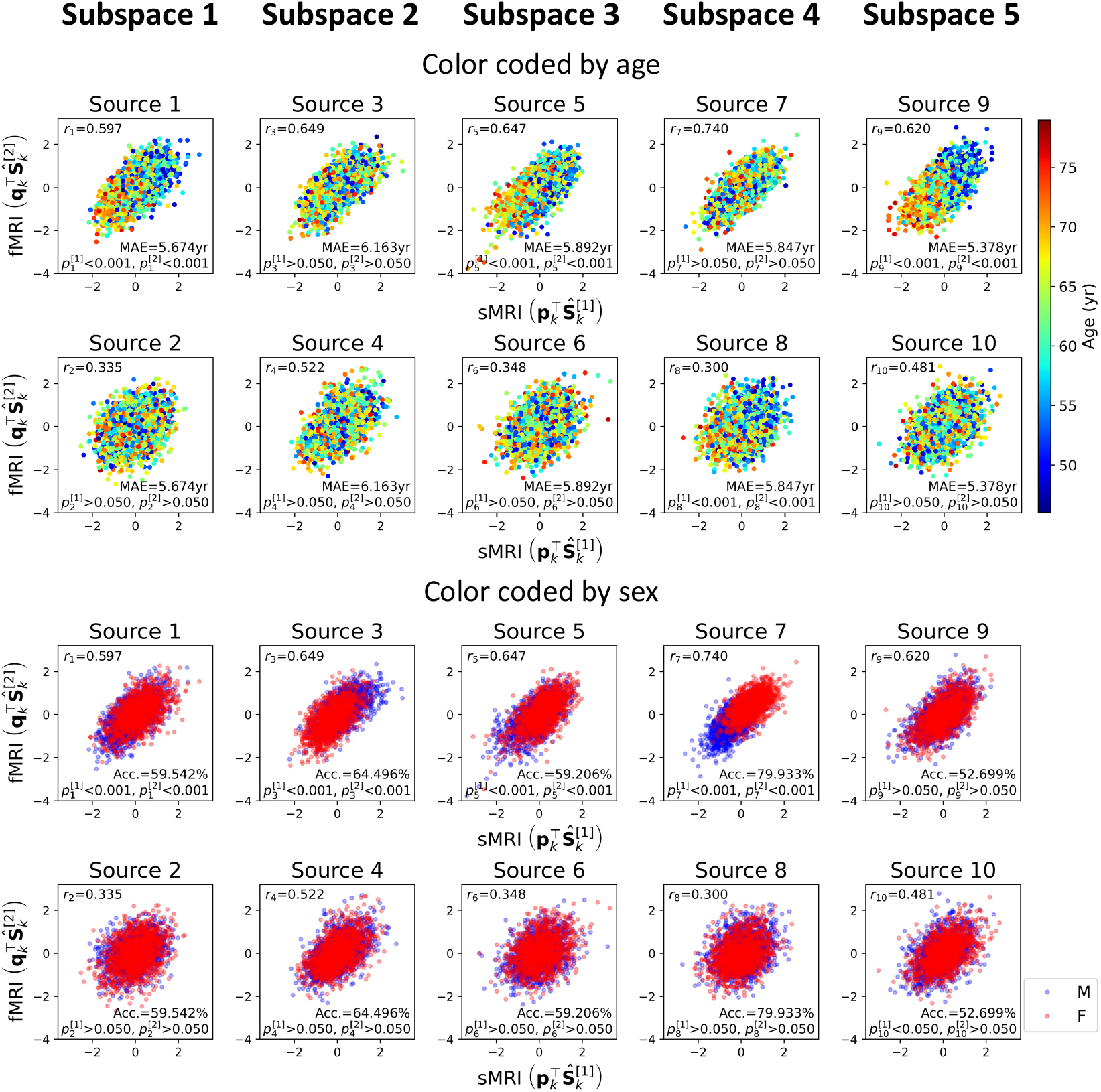
UKB neuroimaging data: Post-CCA sources from MSIVA *S*_2_ cross-modal subspaces, color coded by age or sex. Rows I and II show the age effect; rows III and IV show the sex effect. The Pearson correlation coefficient (*r*_*c*_, *c* is the source index) indicates the correlation of post-CCA sources between two modalities (high *r*_*c*_ means similar expression profiles, not similar spatial organization). We performed age regression and sex classification, measured by mean absolute error (MAE; lower is better) and balanced accuracy (Acc.; higher is better), to assess the associations between MSIVA linked sources from each cross-modal subspace and phenotype measures (age and sex). Sources from subspace 5 yielded the best performance in age regression (MAE = 5.378 years), while sources from subspace 4 achieved the best performance in sex classification (Acc. = 79.933%). The *p*-value indicates statistical difference between two age or sex groups (*p*^[1]^: sMRI; *p*^[2]^: fMRI; two-sample *t*-test, FDR correction for all subjects). Cross-modal sources 1, 5, 8, and 9 were primarily associated with aging (*p*^[1]^ *<* 0.001, *p*^[2]^ *<* 0.001). Cross-modal sources 1, 3, 5, and 7 were more linked to sex differences (*p*^[1]^ *<* 0.001, *p*^[2]^ *<* 0.001).

**Figure 8:**
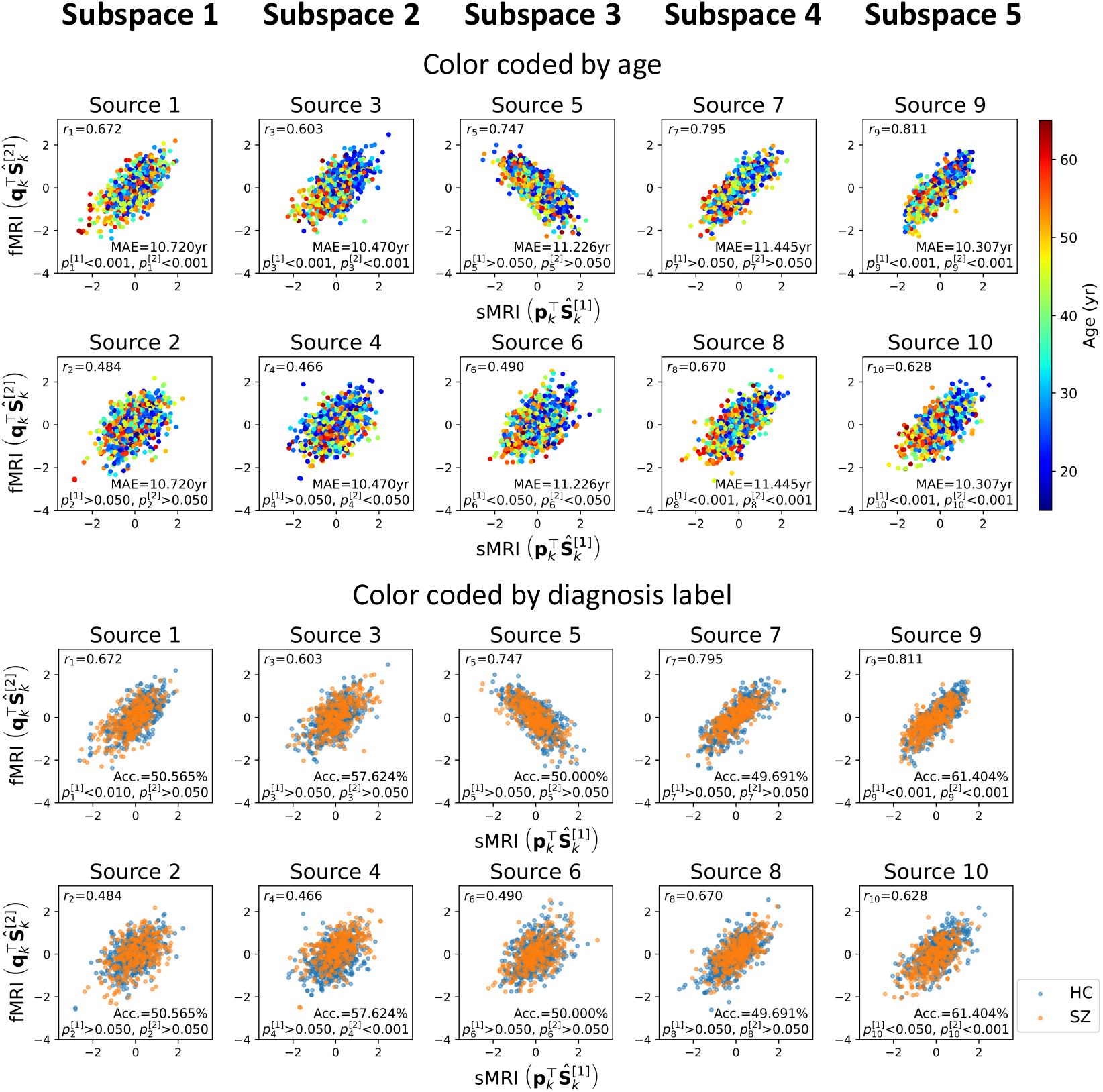
Patient neuroimaging data: Post-CCA sources from MSIVA *S*_2_ cross-modal subspaces, color coded by age or diagnosis labels. Rows I and II show the age effect; rows III and IV show the SZ effect. The Pearson correlation coefficient (*r*_*c*_, *c* is the source index) shows the correlation of post-CCA sources between two modalities (high *r*_*c*_ means similar expression profiles, not similar spatial organization). We performed age regression and diagnosis classification, measured by mean absolute error (MAE; lower is better) and balanced accuracy (Acc.; higher is better), to assess the associations between MSIVA linked sources from each cross-modal subspace and phenotype measures. Sources from subspace 5 yielded the best age regression and diagnosis classification performance (MAE = 10.307 years; Acc. = 61.404%), whereas subspace 2 performed similarly (MAE = 10.470 years; Acc. = 57.624%). The *p*-value indicates statistical difference between two age (younger versus older) or diagnosis (HC versus SZ) groups (*p*^[1]^: sMRI; *p*^[2]^: fMRI; two-sample *t*-test, FDR correction for all subjects). Cross-modal sources 1, 3, 8, 9 and 10 were primarily associated with aging (*p*^[1]^ *<* 0.001, *p*^[2]^ *<* 0.001). Both sources 9 and 10 in subspace 5 exhibited notable associations with SZ-related effects (*p*^[1]^ *<* 0.05, *p*^[2]^ *<* 0.001).

Next, we utilized a dual-coded visualization (Allen et al., 2012) for the modality- and group-specific geometric median spatial maps of the reconstructed data 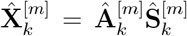 from each representative subspace *k* (Figures 9 and 10). Voxel intensity was mapped to both color hue and opacity. Contours highlighted brain regions with top 15% of voxelwise cross-modal correlations for each group (negative correlations: black; positive correlations: magenta), after eliminating small clusters of voxels using morphological dilation and erosion.

**Figure 9:**
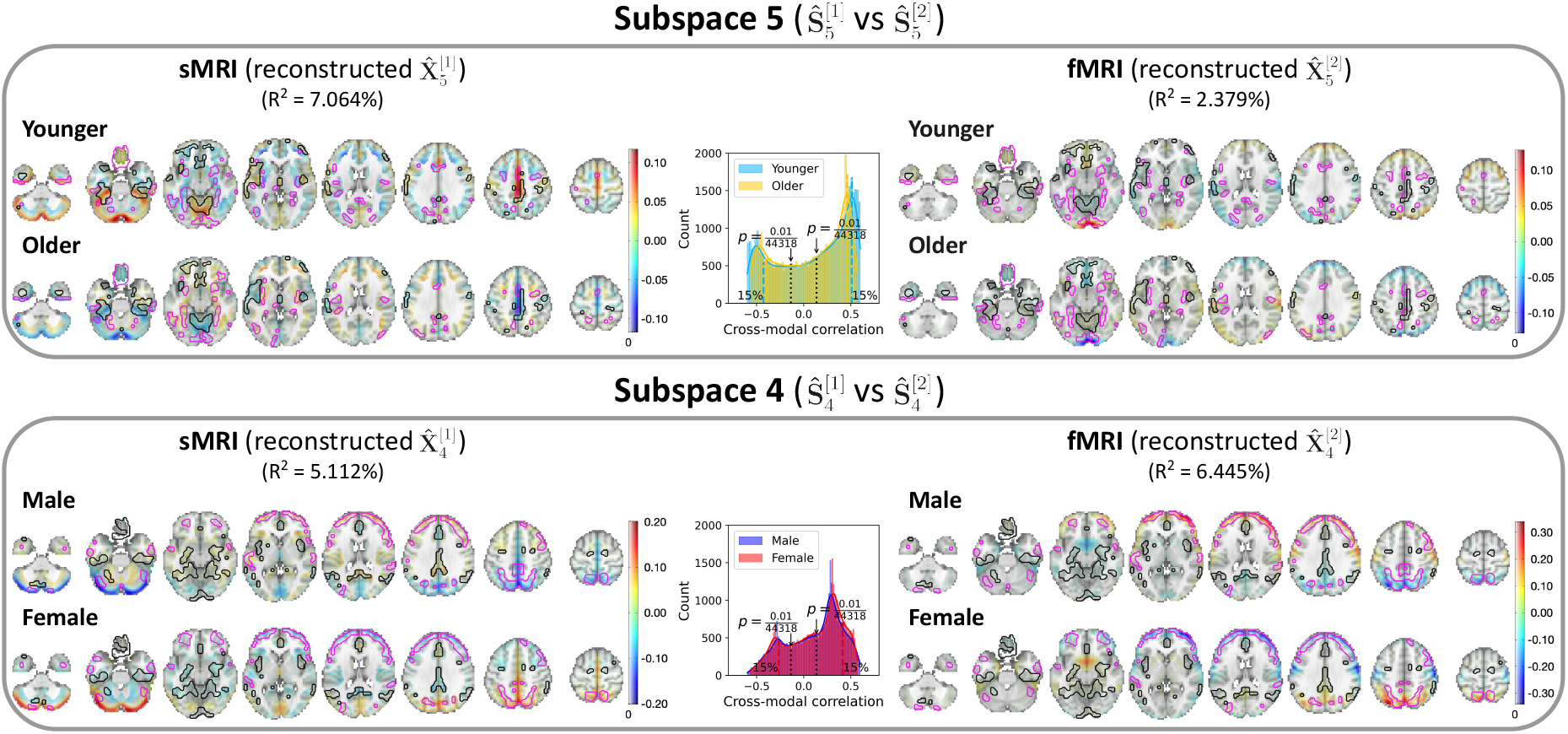
UKB neuroimaging data: Spatial maps of group-specific reconstructed data from MSIVA *S*_2_ sources related to age and sex effects. Axial slices show the geometric median of the reconstructed data 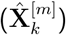 for each modality (sMRI or fMRI) and each group (younger: 46 −62 years, older: 63 −79 years; male or female). Voxel intensity is mapped to both color hue and opacity. Contours highlight brain areas with top 15% of voxelwise cross-modal correlations for each group (negative correlations: black; positive correlations: magenta). Histograms show voxelwise cross-modal correlations for each group (colored dashed lines: top 15% of negative or positive correlations for each group; black dotted lines: 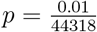, Bonferroni correction for 44318 voxels). The reported *R*^2^ indicates the proportion of variance captured by the subspace in each modality.

**Figure 10:**
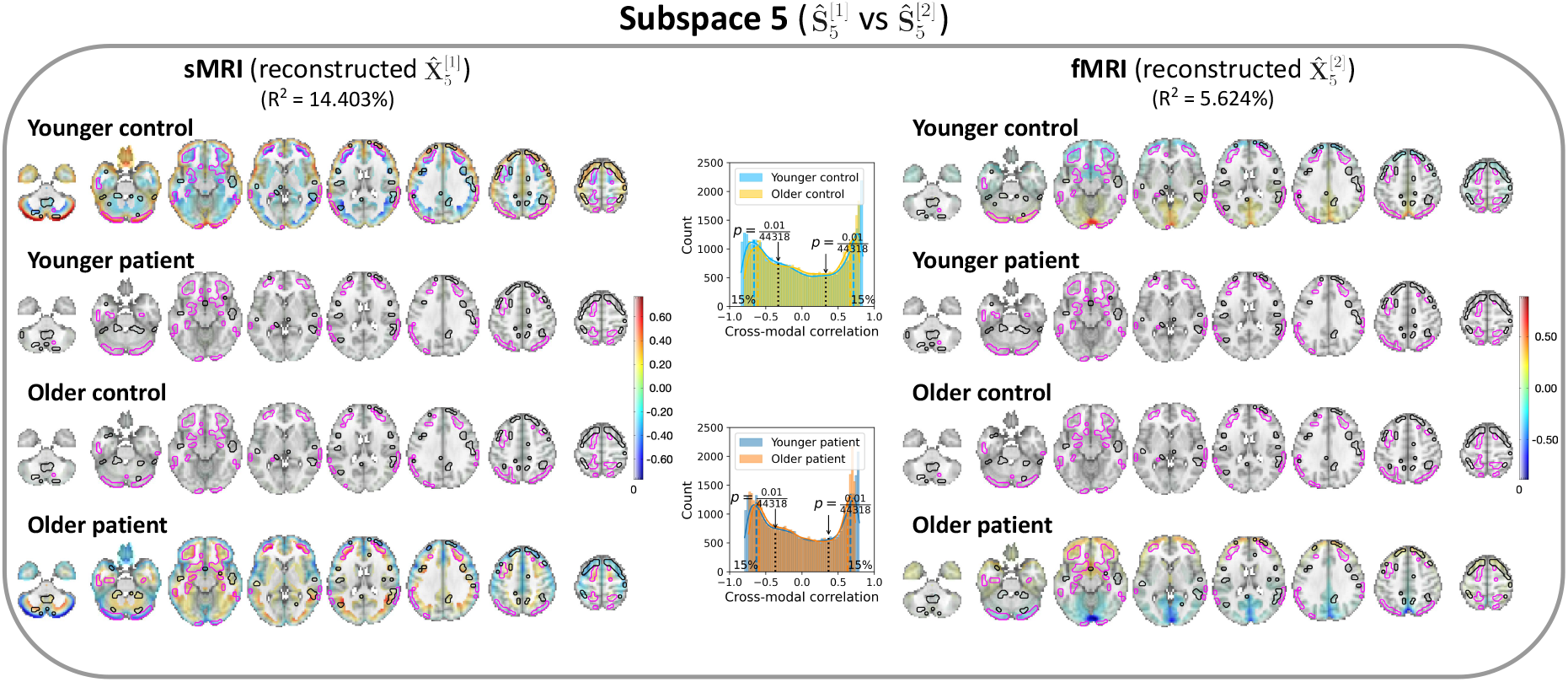
Patient neuroimaging data: Spatial maps of group-specific reconstructed data from MSIVA *S*_2_ sources related to age and SZ interaction effects. Axial slices show the geometric median of the reconstructed data 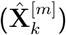 for each modality (sMRI or fMRI) and each group (younger:15− 38 years, older: 39 −65 years; control or patient). Voxel intensity is mapped to both color hue and opacity. Contours highlight brain areas with top 15% of voxelwise cross-modal correlations for each group (negative correlations: black; positive correlations: magenta). Histograms show voxelwise cross-modal correlations for each group (colored dashed lines: top 15% of negative or positive correlations for each group; black dotted lines: 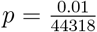, Bonferroni correction for 44318 voxels). The reported *R*^2^ indicates the proportion of variance captured by the subspace in each modality.

In the UKB dataset, source 9 from subspace 5 showed the strongest age effect, while source 7 from subspace 4 showed the strongest sex effect (Figure 9). *Subspace 5* : age effects were observed in the cerebellum, precentral gyrus, cingulate gyrus, and paracingulate gyrus in sMRI; the occipital pole, lateral occipital cortex, superior frontal gyrus, and precuneus in fMRI. In particular, younger subjects (*<* median age in the UKB dataset; 46 − 62 years) showed higher positive voxel intensities in these areas, while older subjects (≥ median age in the UKB dataset; 63 −79 years) showed negative intensities in the same areas. *Subspace 4* : Sex effects were observed in the frontal lobe, occipital lobe, and precuneus in both sMRI and fMRI. Female participants exhibited strong positive intensities in the cerebellum (sMRI), lateral occipital cortex (fMRI), subcallosal area (fMRI), and precuneus cortex (sMRI and fMRI), and negative intensities in the frontal pole and postcentral gyrus (fMRI). Male participants showed the opposite patterns. Spatial maps for the other MSIVA *S*_2_ cross-modal subspaces in the UKB dataset are presented in Appendix I Figure I.1.

In the patient dataset, sources from subspace 5 were significantly associated with different age and diagnosis groups (Figure 10). The younger control participants showed high positive intensities in the cerebellum, temporal pole, and frontal operculum cortex in sMRI; the lingual gyrus, occipital pole, and precuneus cortex in fMRI. They also exhibited negative intensities in the middle temporal gyrus, inferior temporal gyrus, and occipital fusiform gyrus in sMRI. Additionally, both strong positive and negative voxel intensities were observed in the frontal lobe of sMRI. The older patient group (≥ median age in the patient dataset; 39–65 years) showed decreased sMRI intensities in the cerebellum, paracingulate gyrus, insular cortex, frontal pole, middle frontal gyrus, and inferior frontal gyrus, increased sMRI intensities in the middle temporal gyrus and lateral occipital cortex, as well as reduced fMRI intensities in the lingual gyrus, precuneus cortex, and occipital pole, compared to age-matched controls. Spatial maps for the other MSIVA *S*_2_ linked subspaces in the patient dataset are shown in Appendix I Figure I.2.

In addition, the number of voxels with significant cross-modal correlations (*p <* 0.01, Bonferroni correction for 44318 voxels) in older patients diagnosed with SZ (25623) was 18.6% less than their age-matched control subjects (31482) in subspace 5. Particularly, brain areas with reduced structure-function agreement included the insular cortex, lingual gyrus, occipital pole, inferior frontal gyrus, and paracingulate gyrus. Apart from subspace 5, consistent reductions were observed in the other three linked subspaces (Appendix I Figure I.3 subspaces 1-3), suggesting decreased coupling between brain structure and function for older patients.

### 3.4 Brain-age gap is associated with lifestyle factors and cognitive functions

We performed a two-stage voxelwise brain-age delta analysis using the UKB sources estimated by MSIVA using the optimal subspace structure *S*_2_ (see Appendix B for details). We investigated whether the brainage gap was associated with other phenotype variables by computing the Pearson correlation between ***δ***_2*p*_ and each phenotype variable for each voxel. To examine effects specific to shared multimodal variability, we applied voxelwise singular value decomposition (SVD) to the combined reconstructed data from both modalities (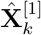and 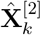) for each of the five cross-modal subspaces. The brainage deltas corresponding to the top SVD-shared voxel-level features of the cross-modal subspaces 2, 4, 5 were significantly associated with various phenotype variables, including time spent watching TV, sleep duration, fluid intelligence, and physical exercise (Figure 11). In particular, predictor 5 (SVD-shared feature from cross-modal subspace 5), which showed the strongest age association (Figure 7 and Appendix B Figure B.1), positively correlated with time spent watching TV and mean time to correctly identify matches (cognitive performance), and negatively correlated with the first principal component of physical exercise variables.

**Figure 11:**
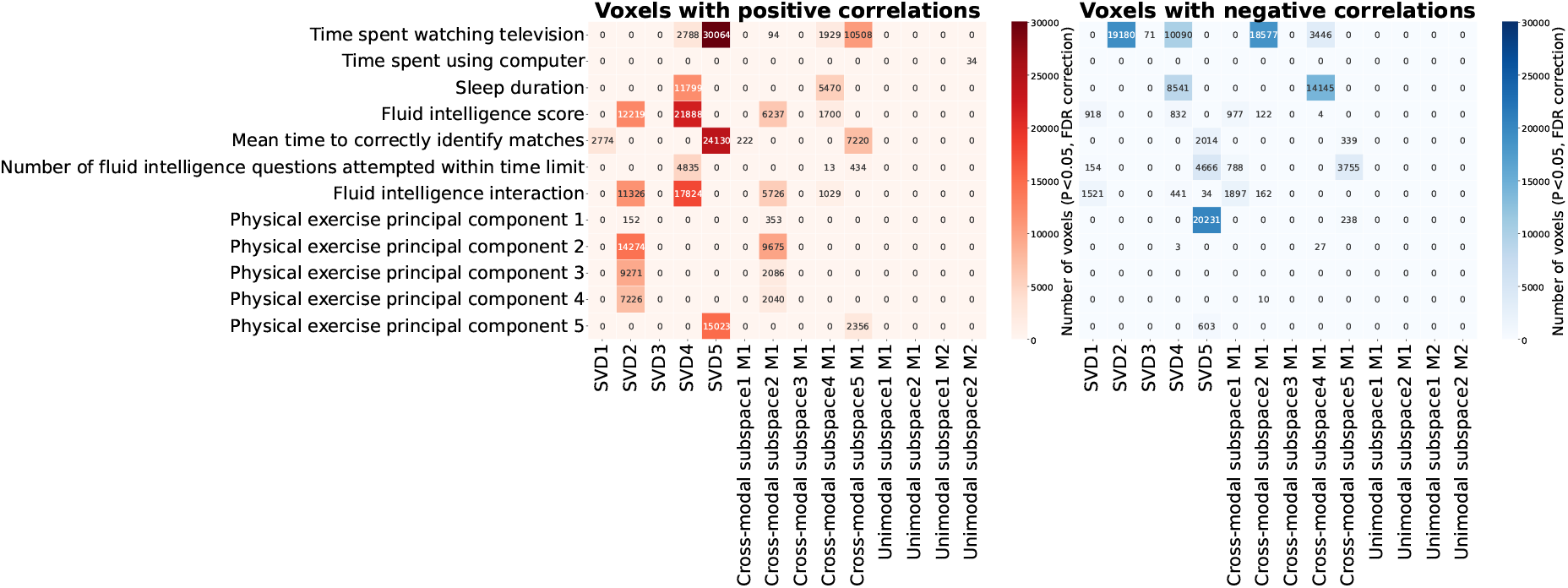
Number of voxels with significant Pearson correlation between corrected brainage delta *δ*_2*p*_ and phenotype variables. Brain-age gap shows significant positive (left) and negative (right) associations with phenotype variables including physical exercise, time spent watching TV, sleep duration, and fluid intelligence (*p <* 0.05, false discovery rate correction for 44318 voxels, 25 phenotype variables, and 14 predictors).

Figure 12 shows the relevant spatial maps of predictor 5 (SVD 5). According to Figure 7, subspace 5 showed the strongest association with the chronological age, which aligned with the strong ***β***_1_ coefficients and *σ*(***δ***_2*p*_) spatial maps from the first step of brain-age delta analysis (Figure 12, panel A, rows I and II). The geometric median of brain-age delta ***δ***_2*p*_ was slightly negative (Figure 12, panel A, row III), indicating that biological age is slightly lower than chronological age (the brain appears younger). We also present spatial maps for three phenotype variables that exhibited strong associations with ***δ***_2*p*_: time spent watching TV, mean time to correctly identify matches, and the first principal component of physical exercise variables (Figure 12, panel B). Particularly, significant effects were observed in the cerebellum, postcentral gyrus, cingulate gyrus, precuneus cortex, occipital lobe, and caudate nucleus for time to watch TV; the frontal pole, precentral gyrus, and insular cortex for time to identify matches; the cerebellum, occipital fusiform gyrus, and caudate nucleus for physical exercise measure. If the correlation on the spatial map is negative (as in the first principal component of physical exercise), ***δ***_2*p*_ decreases as the phenotype score increases and the brain appears younger. If it is positive (as in time to watch TV or identify matches), ***δ***_2*p*_ increases as the phenotype score increases and the brain appears older. Therefore, the more physical exercise, the younger the brain looks; the more time spent watching TV or identifying correct matches, the older the brain looks. These findings indicate that increased physical activity and reduced TV time can potentially improve brain health.

**Figure 12:**
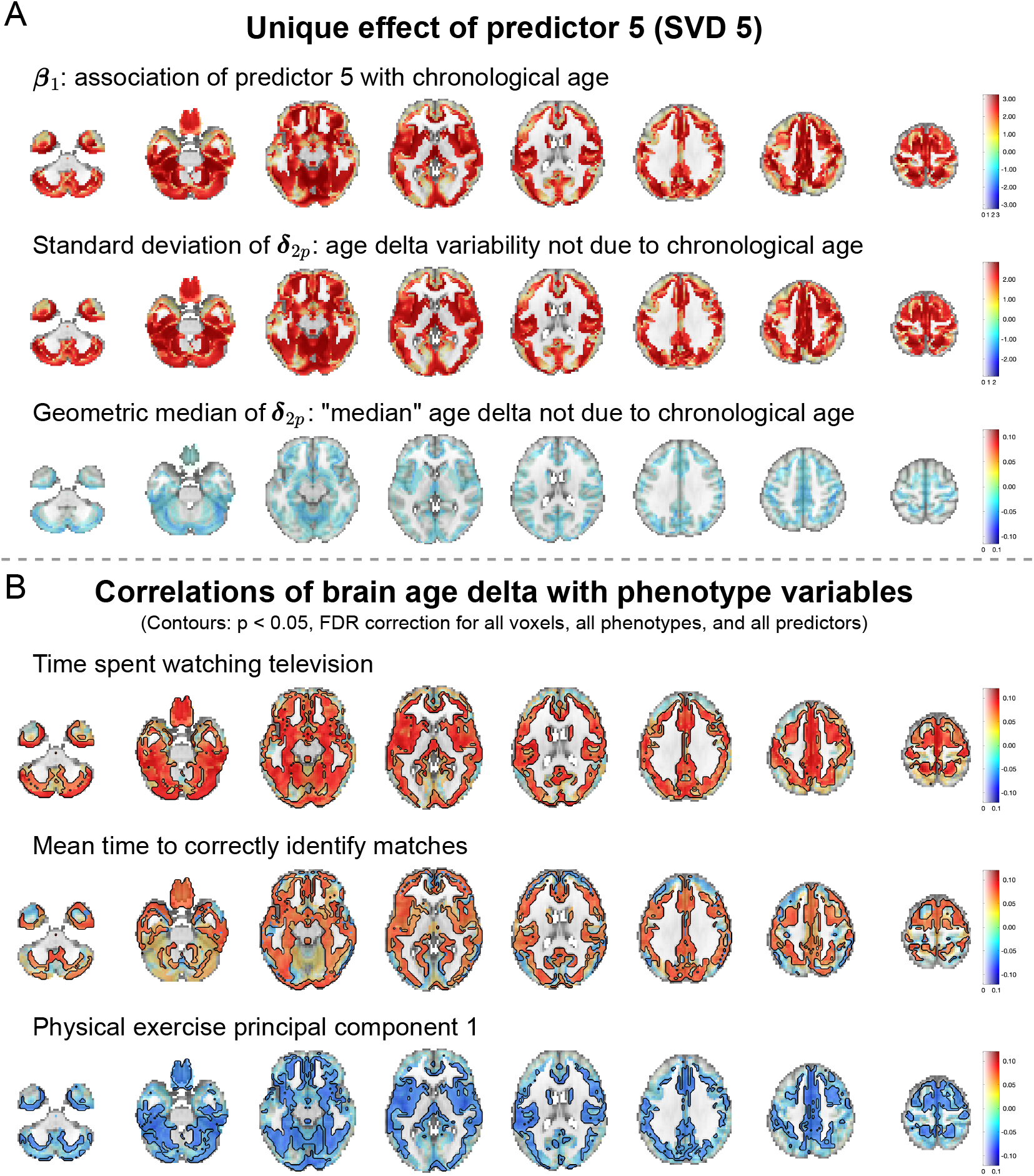
Spatial maps of predictor 5 (SVD 5) from brain-age delta analysis. (A) Spatial maps of ***β***_1_, standard deviation of ***δ***_2*p*_, and geometric median of ***δ***_2*p*_. Voxel value is mapped to both color hue and opacity. (B) Voxelwise correlations between ***δ***_2*p*_ and phenotype variables time spent watching TV, mean time to correctly identify matches, the first principal component of physical exercise variables. The voxelwise correlation is mapped to both color hue and opacity. The contours outline the brain regions where the correlations are significant (*p <* 0.05, false discovery rate correction for 44318 voxels, 25 phenotype variables, and 14 predictors). 14431 voxels overlap within the contours in these three spatial maps.

## 4 Discussion

We present a novel multivariate methodology, Multimodal Subspace Independent Vector Analysis (MSIVA), to capture both cross-modal and unimodal sources. We first showed that MSIVA successfully identified the ground truth when given the correct subspace structure, according to the MISI and interference matrix results, and verified that the correct subspace structures led to the lowest loss values for all synthetic data experiments. We next applied MSIVA to two large multimodal neuroimaging datasets and identified the best-fitted latent subspace structure from five candidates. Among all combinations of different initialization workflows and subspace structures, the MSIVA default initialization method with the subspace structure *S*_2_ output the lowest loss value, thus being considered as the best fit to the latent structure in both neuroimaging datasets. The CCA projections within each cross-modal subspace were strongly associated with age, sex and SZ-related effects, as verified through the phenotype prediction tasks. Moreover, the voxelwise brain-age delta analysis on the UKB dataset identified key non-imaging phenotype variables, including lifestyle factors and cognitive performance, that were significantly correlated with voxel-level brain-age gap.

### Recommendations for model order and subspace structure priors

The success of MSIVA depends on properly identifying the model order and the candidate subspace structures. We propose estimating the number of sources using common information derived from MGPCA, guided by information-theoretic criteria (Y.-O. Li et al., 2007). Since functional imaging sources can form hierarchical clusters ranging from two to four dimensions (Ke et al., 2019; Ma et al., 2010, 2011), we recommend evaluating subspaces with sizes within that range. Future research is encouraged to explore a broader set of subspace structures.

### Evaluation of initialization workflows

We evaluated three initialization workflows that captured different amounts of joint information. Interestingly, MSIVA default initialization and unimodal initialization achieved comparative performance and outperformed a multimodal initialization workflow. One possible explanation is that multimodal initialization may overfit the cross-modal information. Conversely, unimodal initialization does not leverage any cross-modal information, which risks yielding potentially unrecoverable misalignment. MSIVA default initialization, which captures an intermediate level of crossmodal information, appears to strike the best balance among the three initialization workflows. Analyses of within- and cross-modal correlations support this hypothesis (Appendix D). Regardless of the initialization method—MSIVA default initialization or unimodal initialization—MISA effectively aligns the sources during the optimization process.

### Comparison of MSIVA and MMIVA

MSIVA can be viewed as an extension of MMIVA which (1) uses a different default initialization method (MSIVA: MGPCA+ICA initialization; MMIVA: MGPCA+GICA initialization) and (2) allows for arbitrary subspace structures (MSIVA: flexible subspace structures like *S*_1_ − *S*_5_ and more; MMIVA: rigid subspace structures like an identity matrix *S*_5_). To further investigate the relationships between the estimated sources from MSIVA and MMIVA, we compared MSIVA (with the subspace structure *S*_2_) and MMIVA by using MSIVA *S*_2_ sources to predict MMIVA sources, as well as using matched MMIVA sources to predict MSIVA *S*_2_ sources. We found that a pair of MSIVA *S*_2_ sources from each subspace could predict variability in more than two MMIVA sources, while pairs of matched MMIVA sources could also explain variability in more than two MSIVA *S*_2_ sources (Appendix J Figures J.1-J.4). Hence, there was no perfect one-to-one mapping between MSIVA *S*_2_ sources and MMIVA sources. We conclude that MSIVA and MMIVA apportion variability to their sources in different ways. We also note that the mismatch appears to be more pronounced in the patient dataset than in the UKB dataset, which may be related to inherent characteristics of the patient data, such as higher population heterogeneity and smaller sample size. Furthermore, we compared phenotype prediction performance using the MSIVA *S*_2_ post-CCA sources from each cross-modal subspace and the matched pairs of MMIVA sources. MSIVA linked sources from a cross-modal subspace more accurately predicted phenotype measures (age and sex in the UKB dataset and diagnosis in the patient dataset) than pairs of independent MMIVA sources, indicating that higher-dimensional subspaces in MSIVA better capture phenotype-related variability than pairs of linked one-dimensional sources in MMIVA (Appendix J Table J.1).

### Brain regions associated with phenotypes and disorders

MSIVA learns complex statistical relationships across two neuroimaging modalities, gray matter (GM) maps from T1-weighted structural MRI (sMRI) and mean-scaled amplitude of low frequency fluctuations (mALFF) maps from resting-state functional MRI (fMRI), by identifying cross-modal dependence in high-dimensional subspaces. This approach reveals structurally and functionally relevant brain networks and identifies source groups associated with specific phenotypes and disorders.

In the UKB dataset, four cross-modal sources (1, 5, 8, and 9) were significantly associated with age groups, with source 9 from subspace 5 showing the strongest association (Figure 7). Spatial maps reconstructed from subspace 5 revealed that older participants exhibited reduced sMRI intensities in the cerebellum, precentral gyrus, cingulate gyrus, and paracingulate gyrus, as well as reduced fMRI intensities in the occipital pole, precuneus cortex, superior frontal gyrus, and lateral occipital cortex (Figure 9). Several age-related brain regions identified in this study align with findings reported in the previous literature. For example, cerebellar volume has been reported to be associated with age-related decline (Jernigan et al., 2001; Luft et al., 1999; Romero et al., 2021). Hogstrom et al., 2013 has observed strong age effects in the precentral gyrus and weak age effects in the cingulate gyrus using structural brain imaging. In addition, the reductions observed in the visual areas from the mALFF feature maps are consistent with our previous findings using MMIVA (Silva et al., 2024). Prior research on functional networks has also identified significant associations with aging in the occipital lobe (Scheinost et al., 2015).

Regarding sex effects, four sources (1, 3, 5, and 7) showed significant associations with sex, with source 7 from subspace 4 showing the strongest effect (Figure 7). Spatial maps from subspace 4 indicated that males had lower sMRI intensities in the cerebellum and precuneus, lower fMRI intensities in the lateral occipital cortex and precuneus, and higher fMRI intensities in the frontal pole and postcentral gyrus (Figure 10). Previous studies have also reported sex differences in the gray matter volume of the cerebellum (Fan et al., 2010) and the precuneus cortex (Ruigrok et al., 2014), as well as in the frontal and occipital areas using functional measures (Tian et al., 2011). In particular, MMIVA has revealed that females exhibit higher cerebellar sMRI intensities than males (Silva et al., 2024).

In the patient dataset, both sources in subspace 5 showed significant associations with aging and schizophrenia (SZ) (Figure 8). Using reconstructed data from subspace 5 sources, we found that SZ patients, compared with age-matched controls, showed decreased gray matter intensities in the cerebellar, frontal, and insular cortices, and increased intensities in the middle temporal gyrus and lateral occipital cortex. They also exhibited distinct functional reductions in the precuneus, lingual gyrus, and occipital pole (Figure 10). These results align with previous findings of significantly reduced cerebellar gray matter volume in SZ patients (Moberget et al., 2018; Picard et al., 2008), as well as SZ-related alterations in anatomical networks involving the frontal, medial temporal, and insular cortices (Bassett et al., 2008). According to cross-modal correlations, the older patient group shows reduced structure-function coupling in the insular cortex, lingual gyrus, occipital pole, inferior frontal gyrus, and paracingulate gyrus (Appendix I Figure I.3). Similarly, Cocchi et al., 2014 used diffusion MRI and resting-state fMRI to examine structure–function coupling in SZ, reporting reduced white matter integrity and functional connectivity in the frontal, temporal, thalamic, and striatal regions, as well as disrupted functional networks in the parietal, occipital, and temporal cortices where coupling remained intact. Notably, both studies identified frontal and occipital regions, suggesting consistent structure–function alterations in these cortical areas across imaging modalities.

Apart from resting-state data, MSIVA can be applied to task-related or naturalistic data to uncover structural–functional relationships under task conditions or naturalistic stimuli, as well as to other high-dimensional modalities such as genomics, thereby advancing understanding of the molecular mechanisms underlying brain development and disorders.

### Limitations and future directions

A limitation of our current work is the subspace structure used in MSIVA. MSIVA selects the best-fitting subspace structure for the data from a predefined set.However, it is not computationally efficient to exhaustively evaluate the merits of other potential subspace structures. Additionally, we make two assumptions on the subspace structure: the cross-modal subspaces share the same dimensionality per modality, and the unimodal subspaces are all one-dimensional. Yet, it is possible that these assumptions might not represent the true underlying structure of the data. In future work, we plan to apply data-driven subspace structures such as the NeuroMark template (Du et al., 2020; Fu et al., 2024), or learn the underlying subspace structure from the data directly in an unsupervised manner. In this study, we favored 12 latent sources to approximate each data modality in part for computational efficiency during the combinatorial optimization, although 12 sources may be insufficient to recover all multimodal links (Song et al., 2016). Further workflow optimization and *joint* model order estimation methods are needed to effectively determine the sizes and efficiently estimate the alignment of subspaces with higher dimensionality.

We used the loss value to select the optimal subspace structure in the neuroimaging data in the absence of ground-truth information. However, its baseline can vary across initialization methods, thus it may not always serve as a gold standard of goodness of fit. However, when the same initialization workflow is used, loss values demonstrate consistency across test subspace structures, showing significant correlation with MISI values. Therefore, we recommend comparing loss values only under the same initialization. Also, we suggest evaluating performance using multiple metrics in addition to the loss value, such as the cross-modal mean correlation coefficient (CMCC) and the cross-modal minimum distance (CMD), which measure the cross-modal subspace alignment without groud-truth information, exercising caution to not over interpret these as indicators of correct estimation but rather as diagnostics to grossly verify that the results are consistent with the tested subspace structure.

While this study focused on two modalities, MSIVA can be applied to two or more modalities by design. We encourage future work to evaluate its performance on additional data modalities. Another promising future direction is to evaluate alternative initialization methods. For instance, the geometric mean or Log-Euclidean framework—commonly regarded as more principled in applications such as diffusion tensor imaging (DTI) (Arsigny et al., 2006) and electroencephalography (EEG) (Tibermacine et al., 2024)— could serve as valid alternatives to adjust covariance matrices before aggregation in MGPCA. In addition, a key limitation is the linear mixing assumption in MSIVA. MSIVA, like ICA, assumes that each data modality is a linear mixture of latent sources, but the true mixing process in neuroimaging data may be nonlinear, especially considering the multiple physiological effects and nonlinear transformations during fMRI modeling and data preprocessing stages. To address this limitation, we plan to develop *nonlinear* latent variable models that estimate multimodal sources which are nonlinearly mixed.

## 5 Conclusions

Our proposed multivariate methodology MSIVA effectively captures both within- and cross-modal sources, as well as their underlying subspace structure, from multiple synthetic and neuroimaging datasets. According to brain-phenotype modeling, the estimated sources from the MSIVA cross-modal subspaces are strongly associated with phenotype variables including age, sex, and psychosis. Subsequent brain-age delta analysis shows that voxel-wise brain-age gap in the recovered cross-modal subspaces is strongly related to lifestyle and cognitive function measures. Our results support that MSIVA can be applied to uncover linked phenotypic and neuropsychiatric biomarkers of brain structure and function at the voxel level from multimodal neuroimaging data.

## Ethics statement

The authors have no conflicts of interest to declare. This study used the UK Biobank Resource under Application Number 34175. Ethical approval was not required, as confirmed by the license attached to the open access data. Large language models such as Claude and ChatGPT were used to correct grammar mistakes at the sentence level.

## Data and code availability

The UK Biobank dataset can be accessed at https://www.ukbiobank.ac.uk/. The BSNIP and MPRC datasets are available through the NIMH Data Archive (NDA) https://nda.nih.gov/. The COBRE dataset is available from the Collaborative Informatics and Neuroimaging Suite (COINS) https://coins.trendscenter.org/. The FBIRN phase III dataset cannot be shared directly due to the Institutional Review Board (IRB) restrictions. Individuals interested in requesting access can contact Vince D. Calhoun.

Analysis and visualization code for this study is publicly available at https://github.com/trendscenter/MSIVA.git. Code for brain-age delta analysis is adapted from https://www.fmrib.ox.ac.uk/datasets/BrainAgeDelta/. Code for dual-coded images is adapted from https://trendscenter.org/x/datavis/.

## Author contributions

**Xinhui Li**: Conceptualization; formal analysis; investigation; methodology; software; validation; visualization; writing - original draft; writing - review and editing. **Peter Kochunov**: Data curation; writing - reviewing and editing. **Tulay Adali**: Funding acquisition; investigation; writing - reviewing and editing. **Rogers F. Silva**: Conceptualization; formal analysis; investigation; methodology; project administration; software; supervision; writing - review and editing. **Vince D. Calhoun**: Conceptualization; funding acquisition; investigation; project administration; resources; supervision; writing - review & editing.

## Acknowledgments

We acknowledge the FBIRN team who coordinated and performed the data acquisition, including Adrian Preda, Aysenil Belger, Bryon A. Mueller, Daniel H. Mathalon, Daniel S. O’Leary, Jessica A. Turner, Juan R. Bustillo, Judith M. Ford, Kelvin O. Lim, Steven G. Potkin, and Theo G.M. van Erp.

## Appendix A Data acquisition and preprocessing

### A.1 UK Biobank dataset

#### A.1.1 Acquisition parameters

T1-weighted structural MRI (sMRI) images were acquired using a 3D MPRAGE sequence with the following parameters: repetition time (TR) = 2000ms, inversion time (TI) = 880ms, in-plane acceleration factor = 2, voxel size = 1×1×1mm^3^, acquisition matrix = 208×256×256. Resting-state functional MRI (fMRI) were acquired with the following parameters: TR = 735ms, echo time (TE) = 39ms, multiband factor = 8, in-plane acceleration factor = 1, flip angle = 52^°^, voxel size = 2.4×2.4×2.4mm^3^, acquisition matrix = 88 × 88 × 64.

##### A.1.2 Preprocessing steps

For sMRI preprocessing, we performed tissue segmentation and normalization to the Montreal Neurological Institute (MNI) template using the statistical parametric mapping toolbox (SPM12, http://www.fil.ion.ucl.ac.uk/spm/) (Ashburner et al., 2014), leading to gray matter (GM), white matter (WM), and cerebrospinal fluid (CSF) tissue probability maps. Next, the normalized GM tissue probability maps were spatially smoothed using a Gaussian kernel with a full width at half maximum (FWHM) = 10mm. The smoothed images were then resampled to 3 × 3 × 3mm^3^. We next defined a group mask for GM voxels. Specifically, an average GM tissue probability map from all subjects was obtained from the normalized GM tissue probability maps at 1 × 1 × 1mm^3^ resolution. This group-average GM map was binarized at a threshold of 0.2 and resampled to 3 × 3 × 3mm^3^ resolution, resulting in 44318 voxels.

For fMRI preprocessing, we utilized the distortion corrected, FIX-denoised (Griffanti et al., 2014), normalized fMRI data from the UK Biobank data resource to compute subject-specific amplitude of low frequency fluctuations (ALFF) maps, defined as the area under the low frequency band [0.01 − 0.08 Hz] power spectrum of each voxel time course in each scan. We then calculated a mean-scaled ALFF (mALFF) map for each subject, which is the subject-specific ALFF map divided by its global mean ALFF value for greater test-retest reliability (Zhao et al., 2018). The mALFF maps were smoothed using a 6mm FWHM Gaussian filter and resampled to 3 × 3 × 3mm^3^ isotropic voxels. We applied the same group-average GM mask for the mALFF maps, resulting in 44318 voxels.

### A.2 Patient datasets

#### A.2.1 Acquisition parameters

##### BSNIP

We used the BSNIP dataset collected at two sites: (1) Baltimore with a 3-Tesla Siemens Trio Tim scanner and (2) Hartford with a 3-Tesla Siemens Allegra scanner. Isotropic T1-weighted MPRAGE scans were acquired using the following parameters: TR = 6.7ms, TE = 3.1ms, flip angle = 8^°^, matrix size = 256 × 240, total scan time = 10 : 52.6min, 170 sagittal slices, slice thickness = 1mm, voxel size = 1 × 1 × 1.2mm^3^ (Giakoumatos et al., 2015). Resting-state fMRI scans were obtained with the following parameters: (1) Baltimore, TR = 2210ms, TE = 30ms, flip angle = 70^°^, number of slices = 36, voxel size = 3.4 × 3.4 × 4mm^3^, and 140 time points; (2) Hartford, TR = 1500ms, TE = 27ms, flip angle = 70^°^, number of slices = 29, voxel size = 3.4 × 3.4 × 5mm^3^, and 210 time points.

##### COBRE

The COBRE dataset was collected at a single site using a 3-Tesla Siemens Tim Trio scanner. A high-resolution T1-weighted multi-echo MPRAGE sequence was used with the following parameters: TR = 2530ms, TE = [1.64, 3.5, 5.36, 7.22, 9.08]ms, TI = 900ms, flip angle = 7^°^, acquisition matrix = 256 × 256 × 176, voxel size = 1 × 1 × 1mm^3^, number of echos = 5, pixel bandwidth = 650Hz, total scan time = 6min. Resting-state fMRI scans were collected with a standard single-shot full k-space echo-planar imaging (EPI) sequence: TR = 2000ms, TE = 29ms, voxel size = 3.75 × 3.75 × 4.55mm^3^, slice gap = 1.05mm, flip angle = 75^°^, number of slices = 32, field of view (FOV) = 240 × 240mm^2^, matrix size = 64 × 64, and 149 volumes. See https://fcon_1000.projects.nitrc.org/indi/retro/cobre.html for more details.

##### FBIRN

The FBIRN phase III dataset was collected from seven sites. Out of seven sites, six sites used 3-Tesla Siemens Tim Trio scanners and one site used a 3-Tesla General Electric (GE) Discovery MR750 scanner. A high-resolution Siemens MPRAGE sequence was acquired with the following parameters: TR/TE/TI = 2300*/*2.94*/*1100ms, flip angle = 9^°^, acquisition matrix = 256 × 256 × 160. Likewise, a GE IR-SPGR sequence was acquired with the following parameters: TR/TE/TI = 5.95*/*1.99*/*45ms, flip angle = 12^°^, acquisition matrix = 256 × 256 × 166, FOV = 220 × 220mm^2^, voxel size = 0.86 ×0.86 × 1.2 mm^3^, collected in the sagittal plane with GRAPPA/ASSET acceleration factor = 2, and NEX = 1 (Qi et al., 2022). The same resting-state fMRI parameters were used across all seven sites: a standard gradient EPI sequence, TR/TE = 2000*/*30ms, voxel size = 3.4375 × 3.4375 × 4mm^3^, slice gap = 1mm, flip angle = 77^°^, FOV = 220 × 220mm^2^, and 162 volumes (Qi et al., 2022).

##### MPRC

The MPRC dataset was collected at three sites, each using a different 3-Tesla Siemens scanner, with a standard EPI sequence. T1-weighted 3D MPRAGE sequence was collected in the saggital plane with voxel size = 1 × 1 × 1mm^3^ using a Siemens Allegra scanner (TE/TR/TI = 4.3*/*2500*/*1000ms, flip angle = 8^°^) or a Siemens Trio scanner (TE/TR/TI = 2.9*/*2300*/*900ms, flip angle = 9^°^) (Schijven et al., 2023). Resting-state fMRI scans were collected using the following scanners and parameters: 3-Tesla Siemens Allegra scanner (TR/TE = 2000*/*27ms, voxel size = 3.44×3.44×4mm^3^, FOV = 220×220mm^2^, and 150 volumes); 3-Tesla Siemens Trio scanner (TR/TE = 2210*/*30ms, voxel size = 3.44×3.44×4mm^3^, FOV = 220×220mm^2^, and 140 volumes); and 3-Tesla Siemens Tim Trio scanner (TR/TE = 2000*/*30ms, voxel size = 1.72 × 1.72 × 4mm^3^, FOV = 220 × 220mm^2^, and 444 volumes) (Qi et al., 2022).

#### A.2.2 Preprocessing steps

All sMRI datasets were preprocessed using SPM12, following the steps described in Qi et al., 2022. Specifically, the data were normalized to the MNI template using unified segmentation, resampled to 3 × 3 × 3mm^3^, and segmented into GM, WM, and CSF using modulated normalization, leading to GM volume maps. These GM volume maps were then smoothed using a 6mm FWHM Gaussian kernel. To ensure proper segmentation for all subjects, outlier detection was performed using spatial Pearson correlation with the template image.

All fMRI datasets underwent the preprocessing steps as outlined in Qi et al., 2022: removal of the initial five scans to eliminate T1 equilibration effects, slice timing correction, realignment, normalization to the EPI template with 3 × 3 × 3mm^3^ resolution, spatial smoothing using a 6mm FWHM Gaussian kernel, regression of nuisance covariates (including six head motion parameters, CSF, WM) and global signal from the voxelwise time course using a general linear model, and computation of the mALFF maps.

## Appendix B Voxelwise brain-age delta analysis on UK Biobank data

We performed a voxelwise brain-age delta analysis using the estimated sources **Ŝ** from MSIVA subspace structure *S*_2_ in the UK Biobank dataset. We describe the steps to construct imaging-derived predictors as follows.

1. Reconstruction. We first reconstructed the modality- and subspace-specific imaging feature 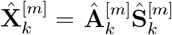 for each of five multimodal subspaces 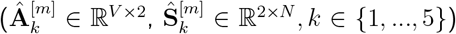 and each of four unimodal subspaces 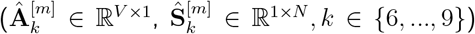, where *k* is the subspace index. Here, subspaces 6 and 7 are used exclusively for sMRI and subspaces 8 and 9 are used exclusively for fMRI.
2. Singular value decomposition (SVD). SVD was applied on cross-modal features to capture the shared information between two modalities. For each voxel *i* in the reconstructed imaging data 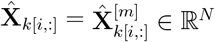 from each of five cross-modal subspaces, we concatenated the two modalities 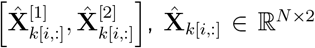, normalized 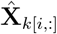 along the rows^8^, and then performed SVD on 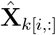, that is, 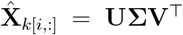. Next, we multiplied 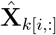 by the first left singular vector 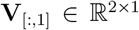 corresponding to the largest singular value *λ*_max_, leading to 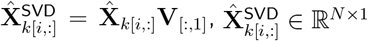 . We then normalized 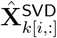 to obtain the normalized 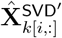.
3. Partialization and normalization. To assess each predictor’s unique variance, we partialized and normalized 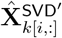 (from five cross-modal subspaces in sMRI) and 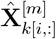 (from four unimodal subspaces in both modalities) to remove SVD-related confounds from 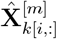, leading to 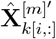. This partialization and normalization procedure follows Smith et al., 2020 to remove confounds from other predictors.
4. Concatenation. We concatenated the SVD results from Step 2 (without partialization or extra normalization) 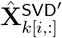 from five cross-modal subspaces, and modality-specific partialized and normalized 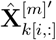 from five cross-modal subspaces and four unimodal subspaces, resulting in 14 predictors in total, 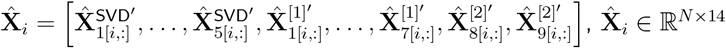.

For each voxel *i*, we performed a two-stage brain age prediction where the first stage estimates the initial delta and the second stage further removes age dependence and other confound factors from the delta (Smith et al., 2019, 2020):

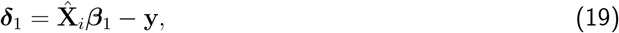

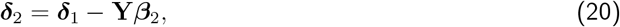

where **y** ∈ ℝ^*N*^ is the chronological age after removing the mean age across subjects. **Y** ∈ ℝ^*N* ×10^ includes the confound variables:

1. the demeaned linear age term,
2. the demeaned quadratic age term after regressing out the linear age effects and normalizing to have the same standard deviation as the linear age term,
3. the demeaned cubic age term after regressing out the linear and quadratic age effects and normalizing to have the same standard deviation as the linear age term,
4. sex,
5. the interaction between sex and each of the three age terms,
6. the framewise displacement variable,
7. the spatial normalization variables from sMRI and fMRI.

Finally, we partialized ***δ***_2_ to remove residual associations, obtaining the partialized brain-age delta, ***δ***_2*p*_.

**Figure B.1:**
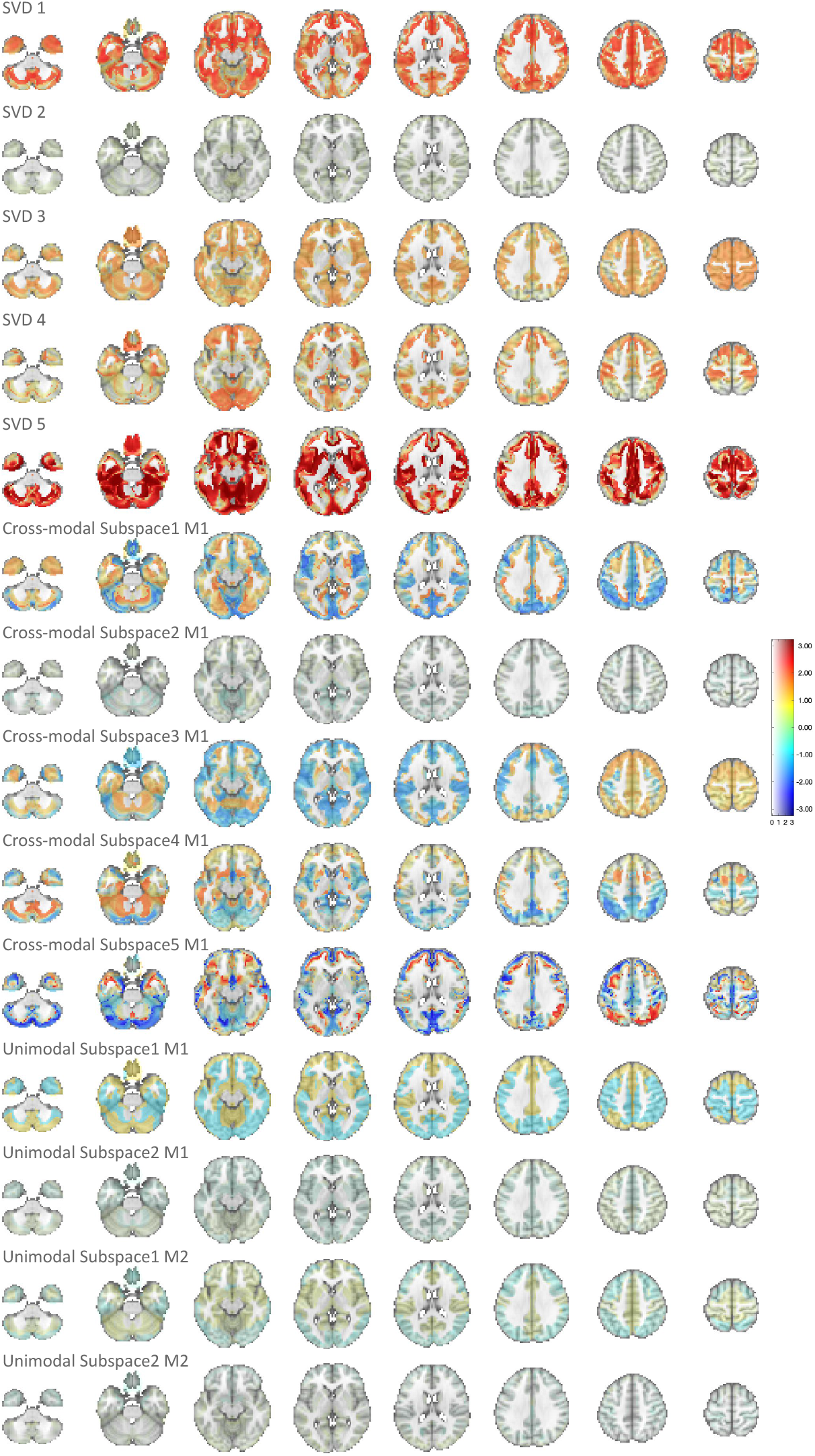
Spatial maps of *β*_1_ in voxelwise brain-age delta analysis. Voxel intensity is mapped to both color hue and opacity. SVD 5 showed the strongest age association among all predictors.

## Appendix C UK Biobank phenotype variables

We used 25 phenotype variables, including lifestyle measures and cognitive test scores, to investigate their associations with brain-age delta. We describe the process of selecting the phenotype variables as follows.

We first excluded variables with extreme values from original 64 non-imaging variables using a two-step approach:

1. We calculated the sum of squared absolute median deviations (*d*) for each variable.
2. We excluded any variable where max(*d*) *>* 100 × mean(*d*), as these extreme outliers could skew statistical analysis.

This initial screening resulted in 54 variables, including age, sex, fluid intelligence, physical activity measures, alcohol intake frequency, cognitive test scores, time spent watching TV, and sleep duration. We further reduced or excluded phenotype variables from these 54 variables as described below:

1. We applied PCA to decompose 28 physical exercise variables into 8 principal components.
2. We removed five age-related variables due to high correlation with other age variables. These variables were “age when attended assessment center”, “age when first sexual intercourse”, “age started wearing glasses”, “years since first sexual intercourse”, and “years since started wearing glasses”.
3. We excluded two variables related to a cognitive test (“time to answer” and “log time to answer”) due to distinct population distributions resulting from two different cognitive tests used during data collection.
4. Finally, we removed the sex variable and another log variable (“log pm score”).

This selection process ultimately yielded 25 variables in total for our analysis, as listed in Table C.1.

**Table C.1:** 54 UK Biobank phenotype variables. 28 physical exercise variables in purple were reduced to 8 principal components by PCA. Variables in red were excluded in brain-age delta analysis. Variables without IDs were created by R.F.S. based on the original variables and not included in the original UK Biobank dataset.

| Variable ID | Variable Name |
| --- | --- |
| f399 2 2 | number of incorrect matches in round |
| f400 2 2 | time to complete round |
| f699 2 0 | length of time at current address |
| f864 2 0 | number of daysweek walked 10 minutes |
| f874 2 0 | duration of walks |
| f884 2 0 | number of daysweek of moderate physical activity 10 minutes |
| f894 2 0 | duration of moderate activity |
| f904 2 0 | number of daysweek of vigorous physical activity 10 minutes |
| f914 2 0 | duration of vigorous activity |
| f943 2 0 | frequency of stair climbing in last 4 weeks |
| f971 2 0 | frequency of walking for pleasure in last 4 weeks |
| f981 2 0 | duration walking for pleasure |
| f991 2 0 | frequency of strenuous sports in last 4 weeks |
| f1001 2 0 | duration of strenuous sports |
| f1011 2 0 | frequency of light diy in last 4 weeks |
| f1021 2 0 | duration of light diy |
| f1050 2 0 | time spend outdoors in summer |
| f1060 2 0 | time spent outdoors in winter |
| f1070 2 0 | time spent watching television tv |
| f1080 2 0 | time spent using computer |
| f1160 2 0 | sleep duration |
| f1438 2 0 | bread intake |
| f1488 2 0 | tea intake |
| f1498 2 0 | coffee intake |
| f1558 2 0 | alcohol intake frequency |
| f2139 2 0 | age first had sexual intercourse |
| f2217 2 0 | age started wearing glasses or contact lenses |
| f2624 2 0 | frequency of heavy diy in last 4 weeks |
| f2634 2 0 | duration of heavy diy |
| f3637 2 0 | frequency of other exercises in last 4 weeks |
| f3647 2 0 | duration of other exercises |
| f4288 2 0 | time to answer |
| f4609 2 0 | longest period of depression |
| f20016 2 0 | fluid intelligence score |
| f20023 2 0 | mean time to correctly identify matches |
| f20128 2 0 | number of fluid intelligence questions attempted within time limit |
| f21003 2 0 | age when attended assessment centre |
| f31 0 0 | sex |
|  | total hours walked 10 minutes |
|  | total hours moderate physical activity 10 minutes |
|  | total hours vigorous physical activity 10 minutes |
|  | total hours of walking for pleasure in last 4 weeks |
|  | total hours of strenuous sports in last 4 weeks |
|  | total hours of other exercises in last 4 weeks |
|  | total hours of light diy in last 4 weeks |
|  | total hours of heavy diy in last 4 weeks |
|  | number of physical activities wrt walking for pleasure |
|  | years since first sexual intercourse |
|  | years since started wearing glasses |
|  | log time to answer |
|  | inverse log duration screen displayed |
|  | inverse log number of attempts |
|  | log pm score |
|  | fluid intelligence interaction |

## Appendix D Comparison of initialization workflows

The choice of weight matrix initialization may affect whether the model can converge to the optimal solution. To better understand the impact of different initialization workflows, we evaluated the initial source estimates obtained after each initialization. Figure D.1 displays the within-modal source correlations (diagonal blocks) and cross-modal source correlations (off-diagonal blocks) following different initialization approaches.

Unimodal initialization applies PCA and ICA to each data modality separately. Since independent sources are estimated without accessing information from other datasets, this method can successfully separate sources within each modality but cannot align sources across modalities, nor group them by subspace within modality (Figure D.1, top row).

In contrast, MSIVA default initialization first applies MGPCA to both datasets simultaneously, followed by ICA on each dimensionally reduced dataset separately. MGPCA identifies orthogonal components using the average covariance matrix across modalities, which preserves cross-modal information during dimensionality reduction. ICA then further separates these MGPCA-derived components into independent sources. This “hybrid” approach effectively separates unimodal sources and identifies substantial cross-modal linkage (evidenced by diagonal structure in cross-modal correlations) for smaller subspace structures (*S*_1_, *S*_2_, and *S*_5_). However, it fails to capture the cross-modal linkage for more complex subspace structures such as *S*_3_ and *S*_4_ (Figure D.1, middle row).

Lastly, multimodal initialization sequentially applies MGPCA and GICA on both datasets. The method demonstrates reasonable results for *S*_5_, which contains one-dimensional sources, but exhibits limitations for other subspace structures containing higher-dimensional subspaces. Specifically, it neither properly separates sources within each modality (high off-diagonal values in within-modal correlations), nor aligns sources between the two modalities, as indicated by the absence of diagonal structure in cross-modal correlations (Figure D.1, bottom row).

While perfect separation and alignment are not expected immediately after initialization—since subsequent combinatorial and numerical optimization steps will further refine these results—achieving a well-balanced combination of unimodal separation and cross-modal alignment during initialization generally increases the likelihood of convergence to the optimal solution.

**Figure D.1:**
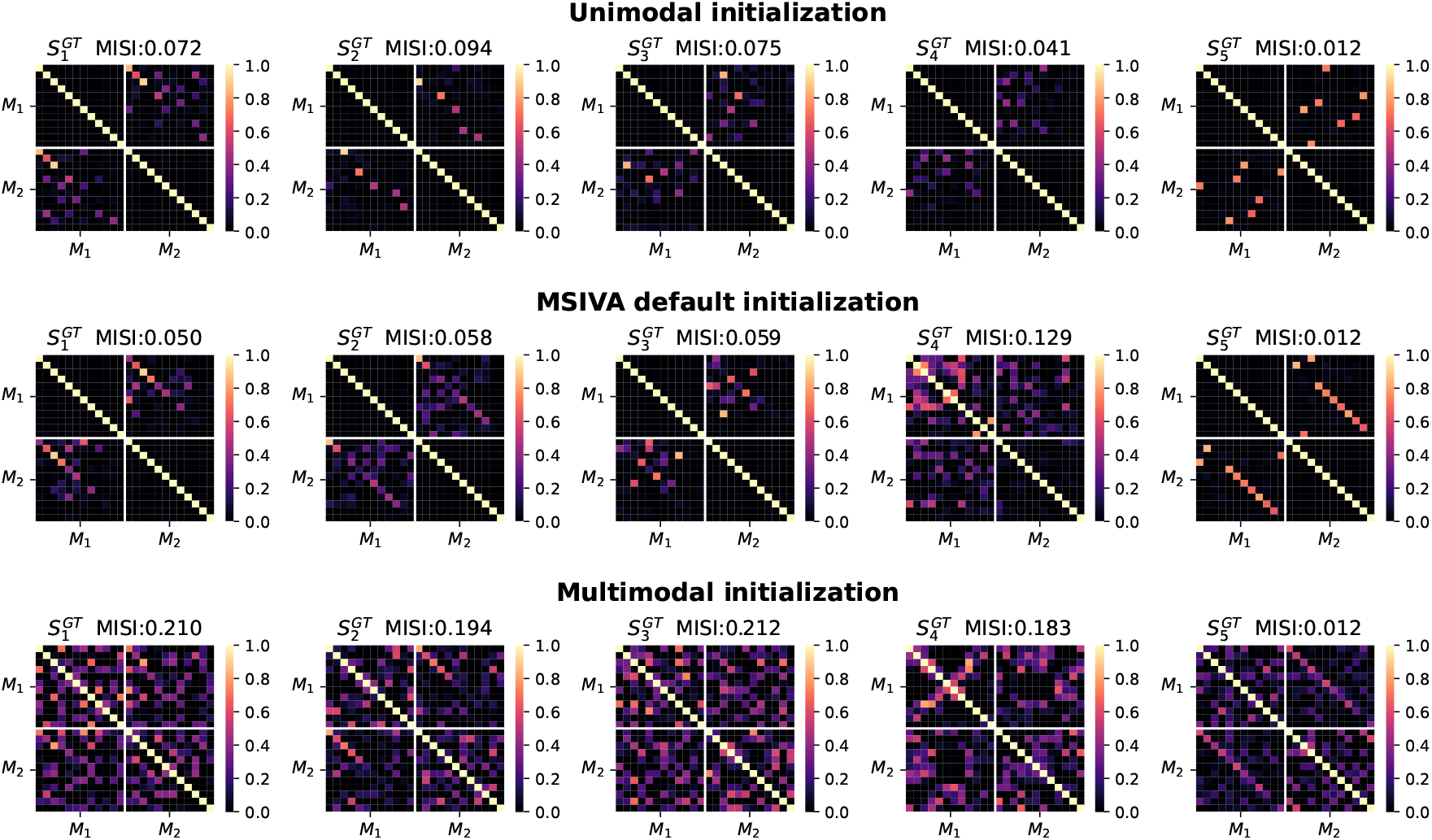
Within-modal and cross-modal source correlations across initialization workflows. From top to bottom, each row shows within- and cross-modal source correlations after unimodal, MSIVA default, and multimodal initialization, respectively. Diagonal blocks represent within-modal correlations; off-diagonal blocks represent cross-modal correlations. MSIVA default initialization achieved the lowest MISI values in four of the five evaluated subspace structures.

## Appendix E Loss curves during numerical optimization

Figure 3 summarizes experiments conducted under 25 conditions and 3 initialization workflows (75 experiments in total). In each experiment, we alternated between combinatorial optimization and numerical optimization 10 times on synthetic data. Each numerical optimization run was capped at 150 iterations. As shown in Figure E.1, the normalized loss curves indicate that most numerical optimization runs converged within 150 iterations (i.e., the loss plateaued), except for multimodal initialization, likely due to suboptimal initialization (Figure D.1). All experiments were executed on an HPC node with two CPUs and 100 GB of memory. Runtime varied substantially across test subspace structures, ranging from 1 minute for 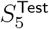 to over 250 minutes for 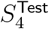. Specifically, 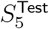 completed in 1 minute, 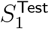 in approximately 20 minutes, 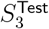 in around 35 minutes, and 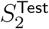 in about 120 minutes. For 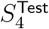, the combinatorial optimization phase was substantially more time-consuming, requiring 253–489 minutes.

Figure E.2 shows the relationships between MISI and other metrics across five subspace structures when the ground-truth and test subspace structures are correctly matched. The loss values were significantly correlated with the MISI values (unimodal initialization: *R*^2^ ≥ 0.95; MSIVA default initialization: *R*^2^ ≥ 0.58), suggesting that loss serves as a valid proxy for MISI in both initialization workflows.

The multimodal mean correlation coefficient (MMCC), which leveraged ground-truth information, showed strong agreement with MISI under both unimodal and MSIVA default initialization workflows (unimodal: *R*^2^ ≥ 0.82; MSIVA: *R*^2^ ≥ 0.79). Similarly, the multimodal minimum distance (MMD) was significantly correlated with MISI for both initialization strategies (unimodal: *R*^2^ ≥ 0.69; MSIVA: *R*^2^ ≥ 0.77). Notably, the minimum MMD value was closer to zero than the minimum value of 1−MMCC, implying that MMD may serve as a more reliable metric than MMCC.

The cross-modal metrics—the cross-modal mean correlation coefficient (CMCC) and the cross-modal minimum distance (CMD)—do not leverage ground-truth information, and were not particularly reliable, as they did not exhibit consistently significant correlations with MISI across the five evaluated subspace structures. Specifically, significant associations were observed only for subspace structures *S*_2_ and *S*_4_ in the unimodal initialization workflow, and for *S*_2_ in the MSIVA default initialization workflow. Among all measures that did not use ground-truth information, only the loss was meaningfully associated with MISI.

**Figure E.1:**
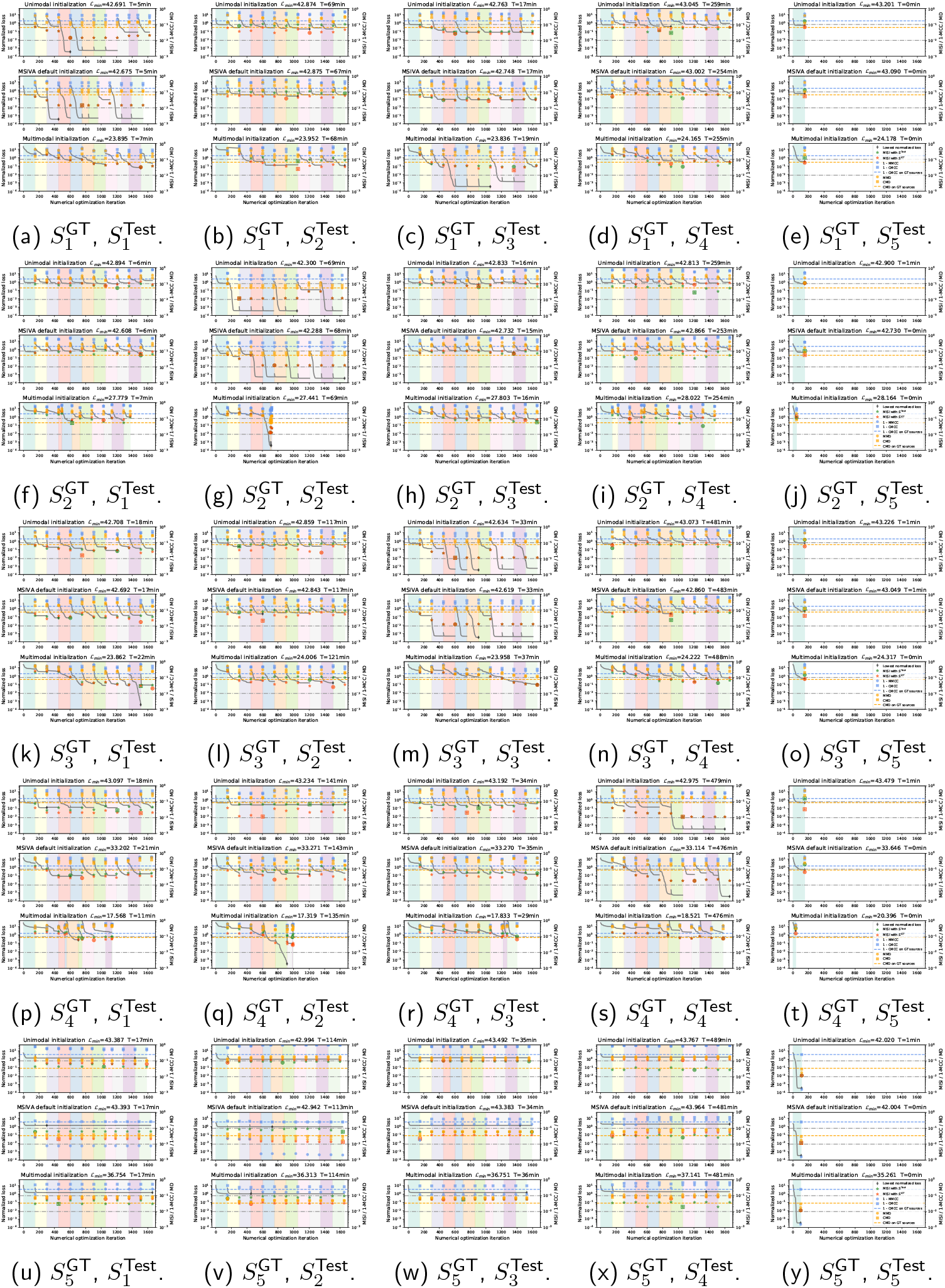
Loss curves during numerical optimization. Panels a, g, m, s, and y (*S*^GT^ = *S*^Test^) confirm that only the unimodal and MSIVA workflows consistently converge to the lowest loss values and detect the correct subspace structures. Vertical colored stripes mark the start and end of numerical optimization runs (≤ 150 iterations each), with combinatorial optimization performed between successive numerical runs. The *left* y-axis shows the normalized loss values after subtracting the minimum loss across the five different test subspace structures *under the same initialization workflow*. The *right* y-axis shows other performance metrics at the end of each numerical optimization run: MISI, 1 − MMCC, 1 − CMCC, MMD, and CMD. The lowest (best) MISI within a workflow is highlighted in a circle (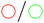), while the lowest MISI across all three workflows is highlighted in a square (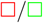).

**Figure E.2:**
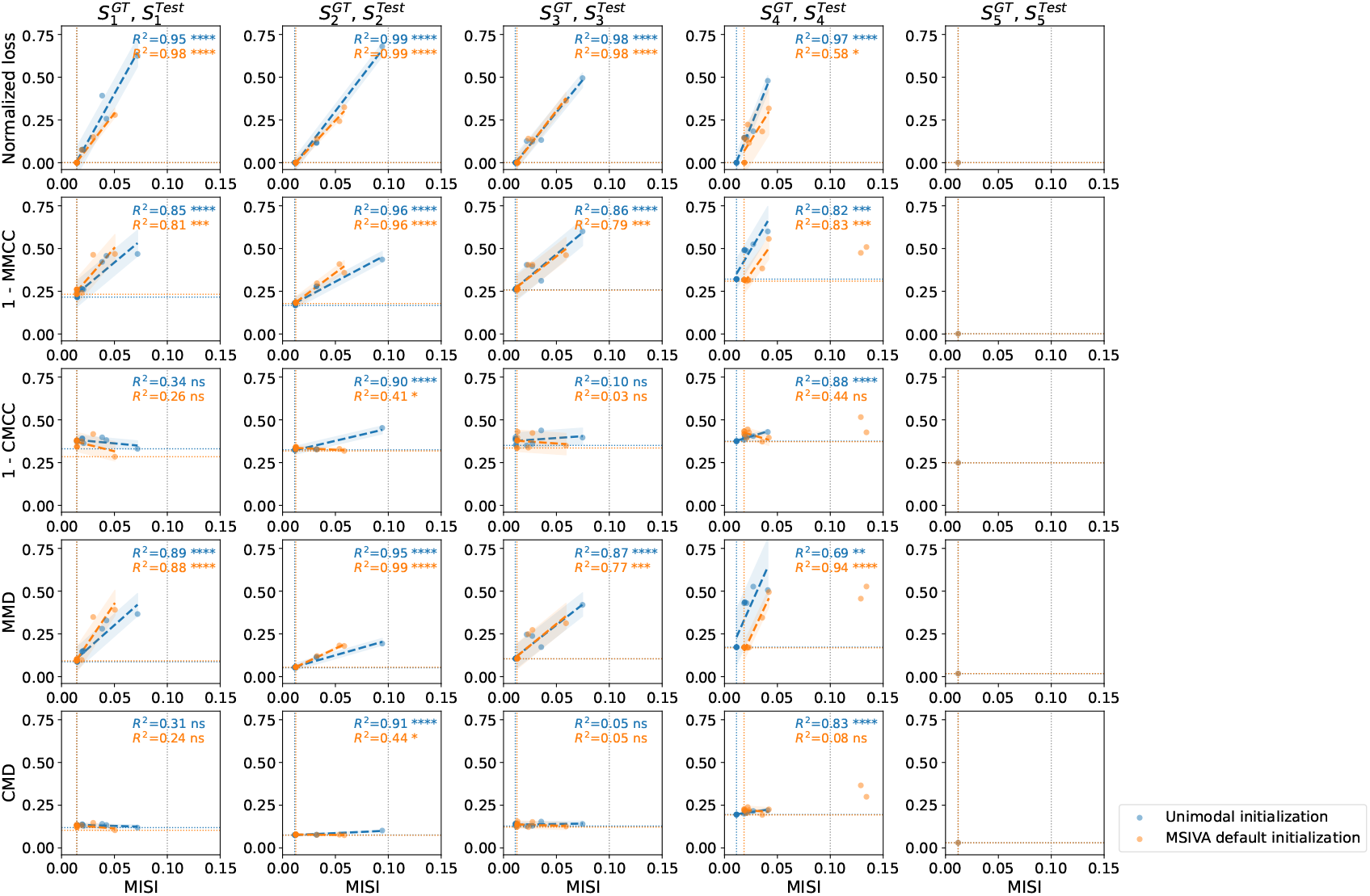
Scatter plots of MISI versus other metrics. Each row presents scatter plots of MISI versus another metric (normalized loss, 1 −MMCC, 1 −CMCC, MMD, or CMD) across the five evaluated subspace structures, under the condition that the test subspace structure matches the ground truth. Each point corresponds to the result from the final iteration of a combinatorial optimization run. In each plot, the blue or orange vertical dotted line indicates the minimum MISI value, and the horizontal dotted line indicates the minimum value of the corresponding comparison metric. The gray vertical line marks the MISI threshold of 0.1, which is commonly used as a heuristic criterion for good source separation (MISI ≤ 0.1). For points with MISI ≤ 0.1, we fit a linear regression line and reported *R*^2^ and *p*-values. The number of asterisks (1, 2, 3, 4) denotes the level of statistical significance in the linear regression analysis (*p <* 0.05, *p <* 0.01, *p <* 0.001, *p <* 0.0001), while “ns” indicates a non-significant result. Normalized loss values were computed by subtracting the minimum loss across the five test subspace structures within the same initialization workflow. Loss values were significantly correlated with MISI, suggesting that loss serves as a valid proxy for MISI. The multimodal metrics, 1 − MMCC and MMD, which leveraged ground-truth information, also showed significant correlations with MISI. In contrast, the cross-modal metrics, 1 − CMCC and CMD, which did not use ground-truth information, were generally not correlated with MISI. Among all measures that did not use ground-truth information, only the loss was meaningfully associated with MISI.

## Appendix F Order selection

In order to select the model order (the number of sources), we evaluated three information-theoretic criteria (Y.-O. Li et al., 2007) based on the singular values of the MGPCA covariance matrix derived from each multimodal neuroimaging dataset: the Akaike Information Criterion (AIC) (Akaike, 1998), the Kullback-Leibler Information Criterion (KIC) (Cavanaugh, 1999), and the Minimum Description Length (MDL) (Rissanen, 1978).

As shown in Figure F.1, the selected orders using AIC, KIC, and MDL are 25, 18, and 10 for the UKB neuroimaging dataset, and 12, 8, and 4 for the patient dataset, respectively. In the patient dataset, the maximum selected order is 12. In the UKB dataset, although AIC and KIC suggest higher orders, we note that the curves are nearly flat as the number of sources is 12 or higher, suggesting that 12 sources already capture a large amount of the information. Considering the increasing computational cost with more sources, we chose 12 sources for both datasets in this study.

**Figure F.1:**
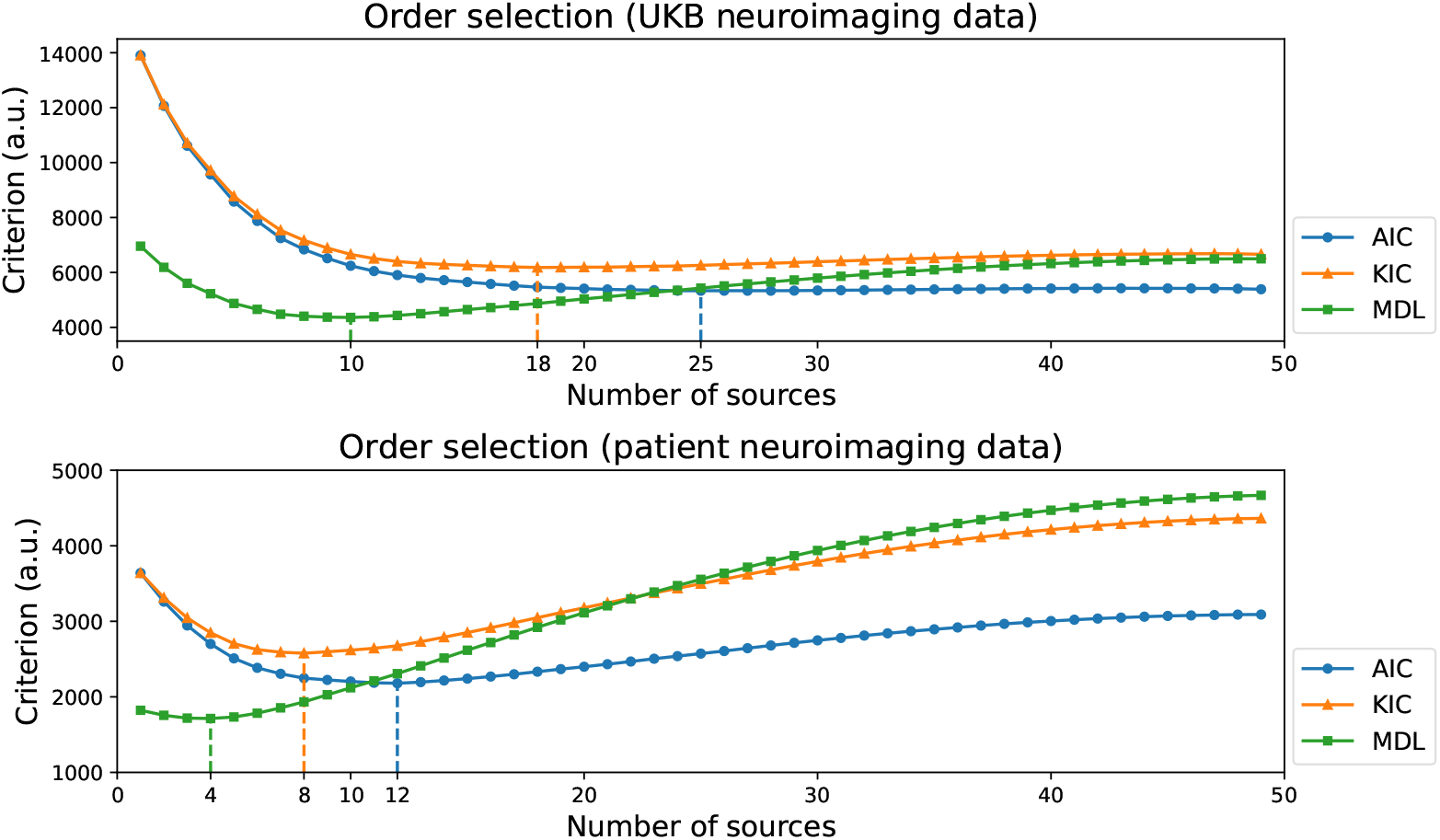
Order selection for multimodal neuroimaging data. The selected orders using AIC, KIC, and MDL are 25, 18, and 10 for the UKB neuroimaging dataset, and 12, 8, and 4 for the patient dataset, respectively. Twelve sources captured the majority of information in the patient dataset and a substantial amount in the UK Biobank dataset. Therefore, we used twelve sources for both datasets in this study.

## Appendix G Nonlinear source dependence in neuroimaging data

Apart from Pearson correlation, we calculated the randomized correlation coefficents (RDCs) (Lopez-Paz et al., 2013) to measure source correlations of random nonlinear copula projections in neuroimaging data. The RDC results (Figures G.1, G.2) were largely consistent with those measured by the linear Pearson correlation coefficient (Figures 5, 6). The low RDC values outside the block subspace structure indicated very weak residual dependence between subspaces, suggesting that different subspaces are nearly independent. The MSIVA default initialization workflow exhibited stronger cross-modal correlations (higher CMCCs and lower CMDs) than the unimodal initialization workflow for all predefined subspace structures (row VI versus row III) in both neuroimaging datasets.

**Figure G.1:**
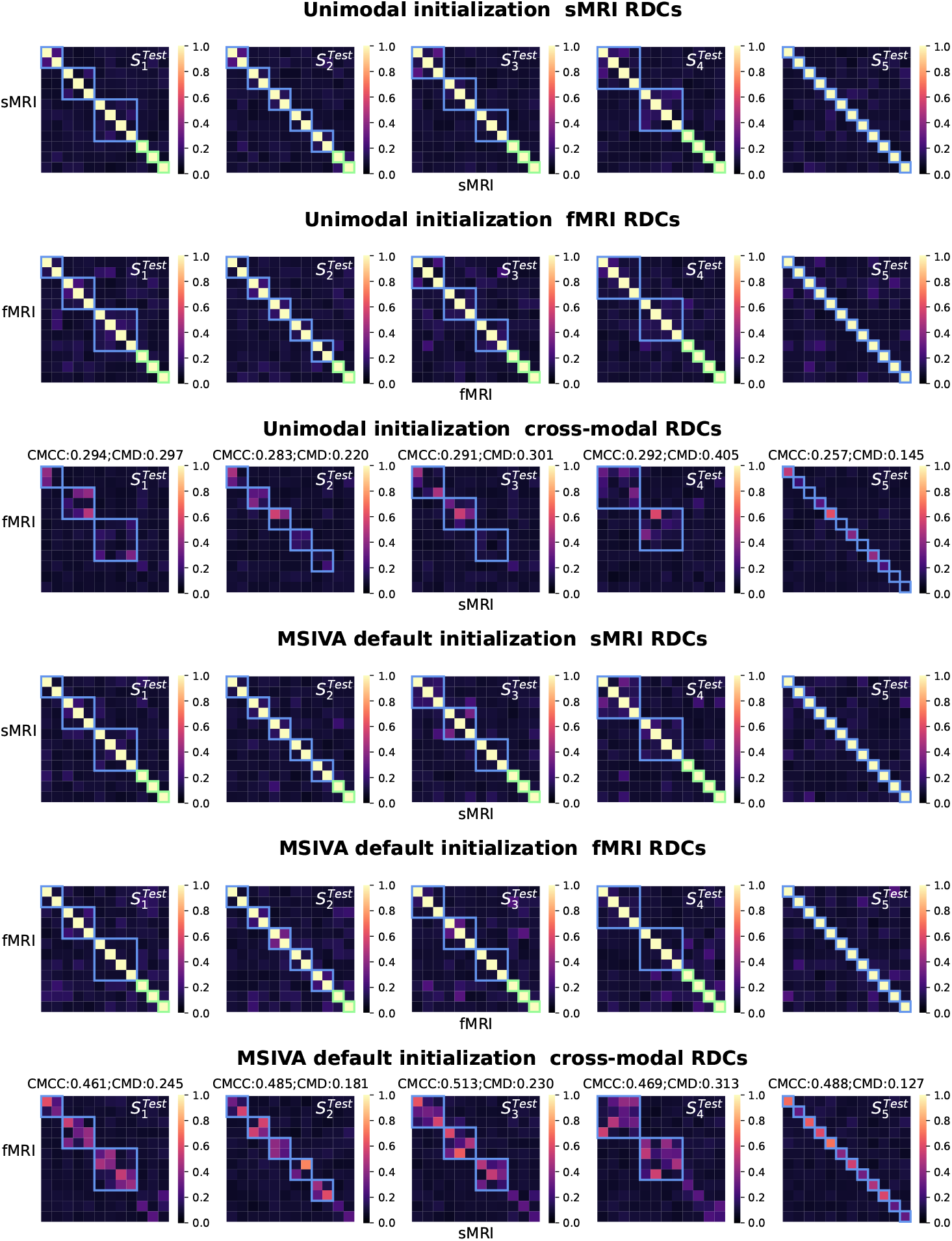
UKB neuroimaging data: Within-modal and cross-modal correlations of random nonlinear copula projections of the recovered sources before applying post-hoc CCA. Crossmodal subspaces are highlighted in blue while unimodal subspaces are highlighted in green. Within-modal self-correlation patterns showed very weak residual dependence between subspaces (rows I-II and IV-V). The MSIVA default initialization workflow exhibited stronger cross-modal correlations (higher CMCCs and lower CMDs) than the unimodal initialization workflow for all predefined subspace structures (row VI versus row III).

**Figure G.2:**
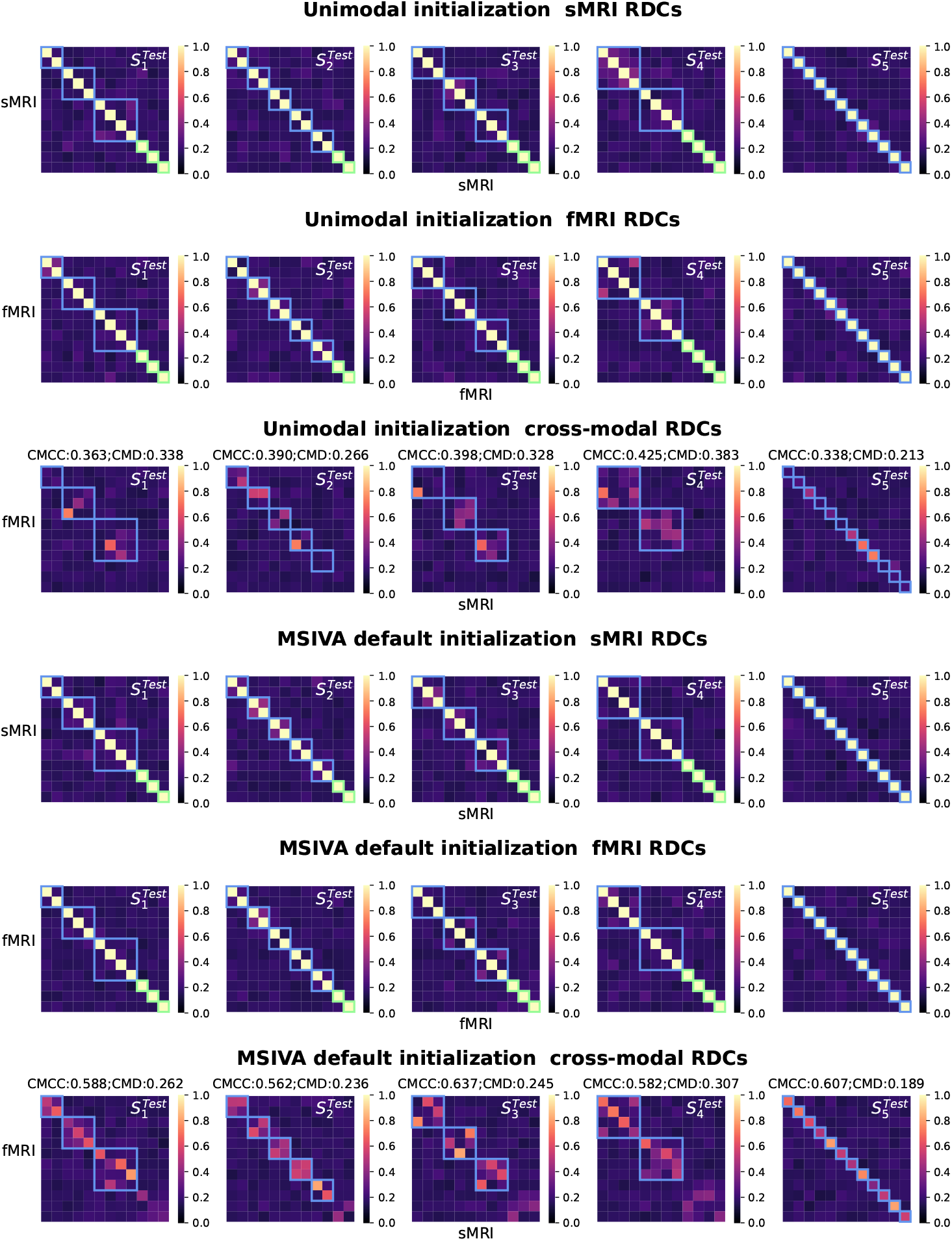
Patient neuroimaging data: Within-modal and cross-modal correlations of random nonlinear copula projections of the recovered sources before applying post-hoc CCA. Crossmodal subspaces are highlighted in blue while unimodal subspaces are highlighted in green. Within-modal self-correlation patterns showed weak residual dependence between subspaces (rows I-II and IV-V). The MSIVA default initialization workflow showed stronger cross-modal correlations (higher CMCCs and lower CMDs) than the unimodal initialization workflow for all predefined subspace structures (row VI versus row III).

## Appendix H Sex effects in patient neuroimaging data

For the patient neuroimaging data, we evaluated sex effects in the post-CCA sources from MSIVA *S*_2_ cross-modal subspaces (Figure H.1). The balanced accuracy for sex prediction was approximately 50% for each source. Moreover, no significant sex effects were observed for any source (*p*^[1]^ *>* 0.05, *p*^[2]^ *>* 0.05 for all sources; two-sample *t*-test with FDR correction for all subjects).

**Figure H.1:**
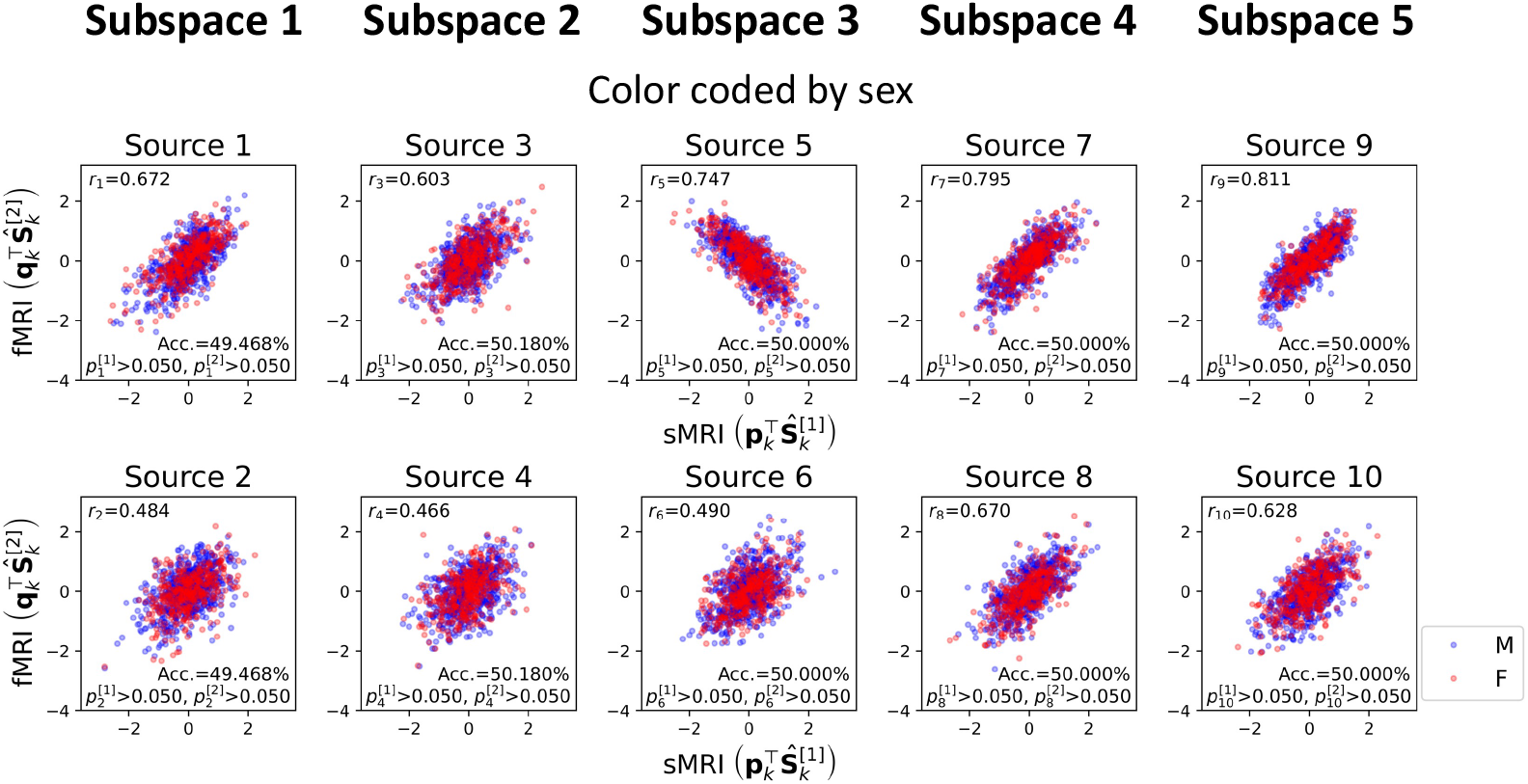
Patient neuroimaging data: Post-CCA sources from MSIVA *S*_2_ cross-modal subspaces, color coded by sex. The Pearson correlation coefficient (*r*) shows the correlation of post-CCA sources between two modalities. The *p*-value shows statistical differences between two sex groups for sMRI and fMRI, respectively (*p*^[1]^: sMRI; *p*^[2]^: fMRI; two-sample *t*-test, FDR correction for all subjects). Sex effects were not significant in the patient neuroimaging dataset.

## Appendix I MSIVA *S*_2_ reconstructed neuroimaging data

Figure I.1 shows spatial maps of group-specific reconstructed data from each of five MSIVA *S*_2_ linked subspaces in the UKB dataset. Similarly, Figure I.2 shows results related to the age and SZ interaction effects in the patient dataset. In each panel, axial slices show the geometric median of the reconstructed subspace data 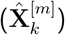 for each modality and each group. Voxel intensity is mapped to both color hue and opacity. Contours highlight brain areas with top 15% of voxelwise cross-modal correlations for each group (negative correlations: black; positive correlations: magenta). Histograms show voxelwise cross-modal correlations for each group (colored dashed lines: top 15% of negative or positive correlations for each group; black dotted lines: 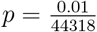, Bonferroni correction for 44318 voxels). The reported *R*^2^ indicates the proportion of variance captured by the subspace in each modality.

Figure I.3 illustrates the number of voxels that show significant cross-modal correlations for age and sex groups in the UKB dataset (rows I and II), and for age and diagnosis groups in the patient dataset (rows III and IV). The number of voxels for older patients diagnosed with SZ is consistently less than that for their age-matched control subjects in four of five subspaces, implying reduced brain structure-function coupling in the older patient group.

**Figure I.1:**
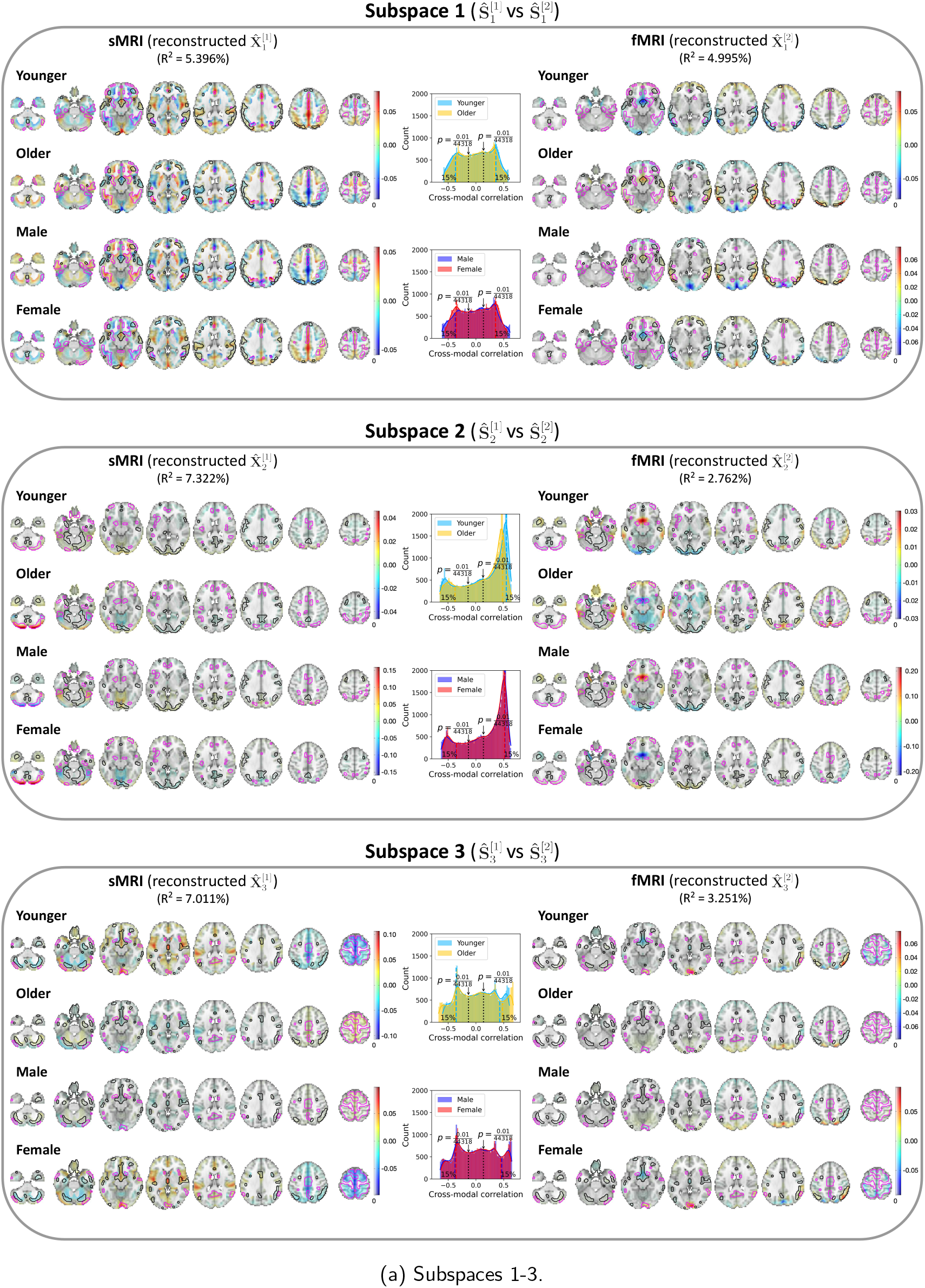
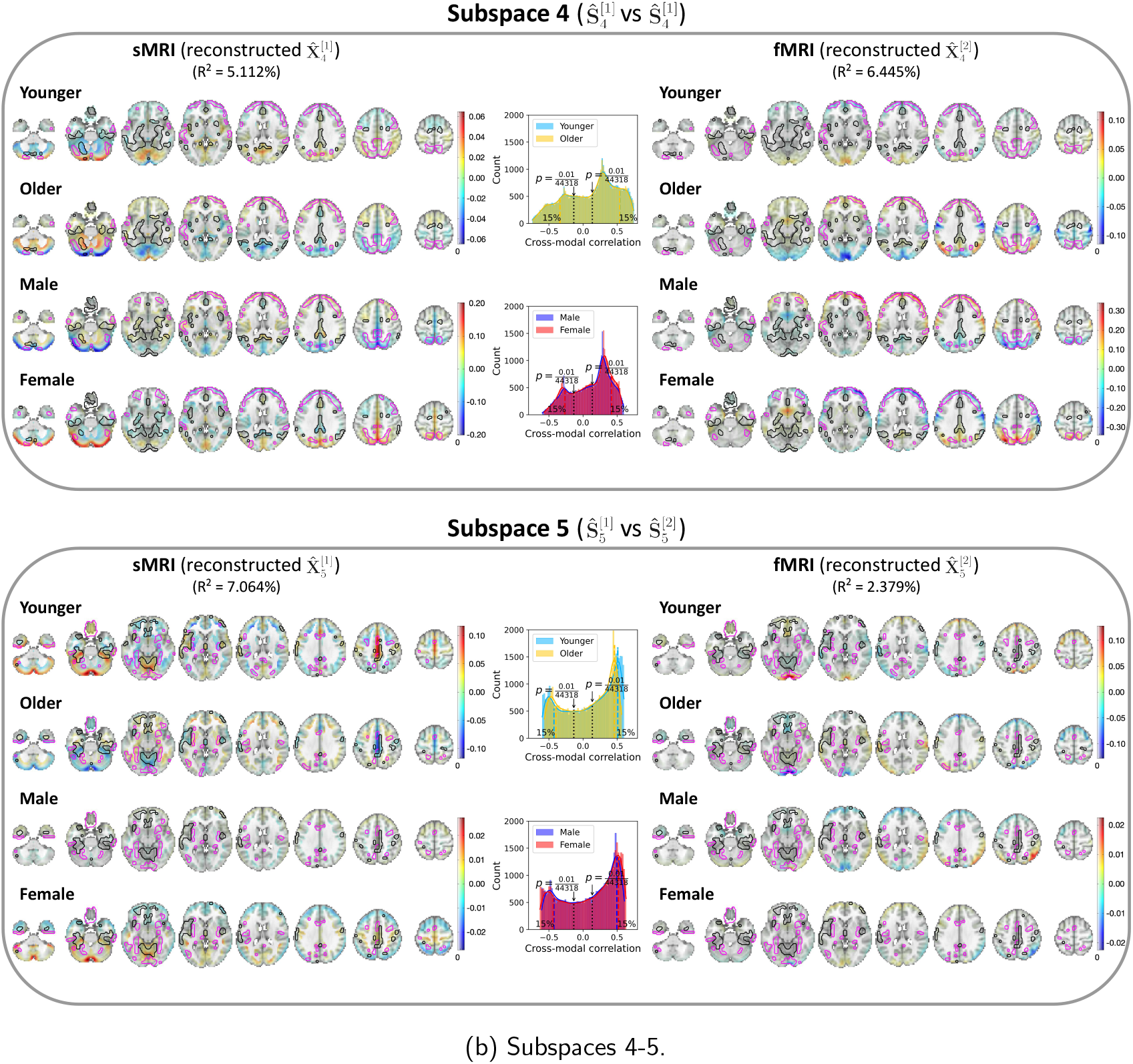
UKB neuroimaging data: Spatial maps of group-specific reconstructed data from MSIVA *S*_2_ sources related to age and sex effects. Axial slices show the geometric median of the reconstructed data 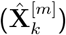 for each modality (sMRI or fMRI) and each group (younger: 46− 62 years, older: 63 −79 years; male or female). Voxel intensity is mapped to both color hue and opacity. Contours highlight brain areas with top 15% of voxelwise cross-modal correlations for each group (negative correlations: black; positive correlations: magenta). Histograms show voxelwise cross-modal correlations for each group (colored dashed lines: top 15% of negative or positive correlations for each group; black dotted lines: 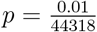, Bonferroni correction for 44318 voxels). The reported *R*^2^ indicates the proportion of variance captured by the subspace in each modality.

**Figure I.2:**
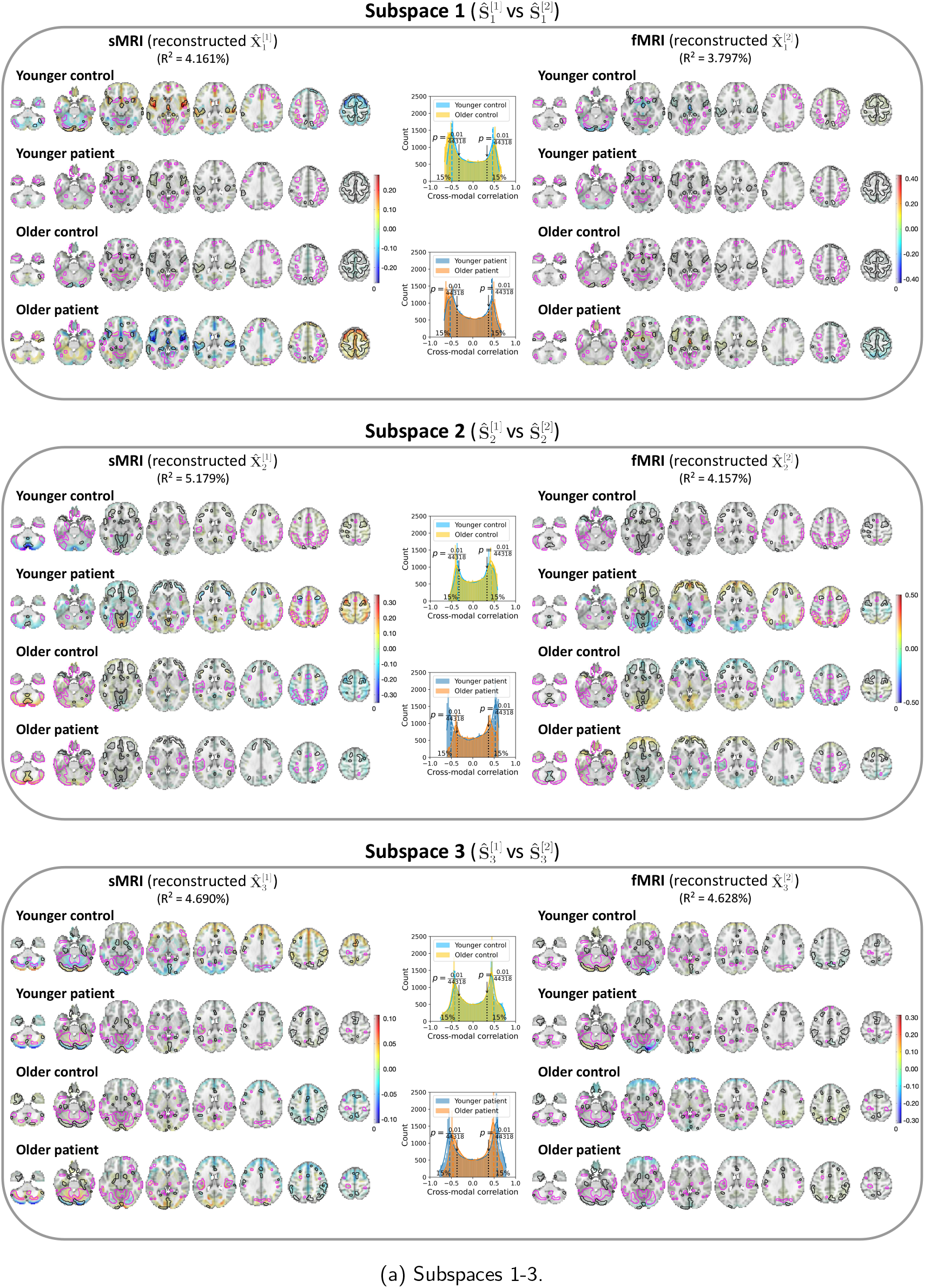
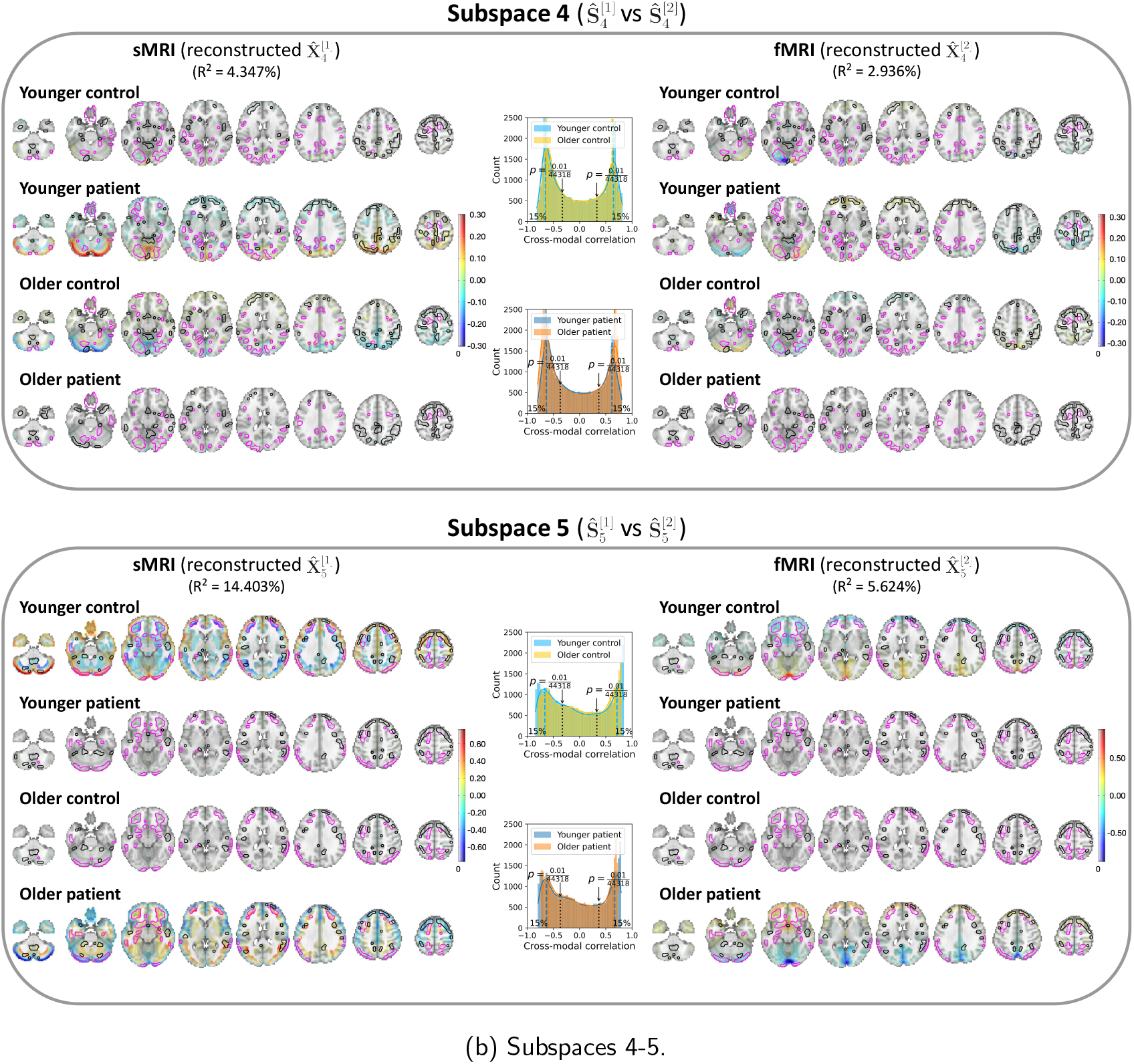
Patient neuroimaging data: Spatial maps of group-specific reconstructed data from MSIVA *S*_2_ sources related to age and SZ interaction effects. Axial slices show the geometric median of the reconstructed data 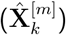 for each modality (sMRI or fMRI) and each group (younger: 15 −38 years, older: 39 −65 years; control or patient). Voxel intensity is mapped to both color hue and opacity. Contours highlight brain areas with top 15% of voxelwise cross-modal correlations for each group (negative correlations: black; positive correlations: magenta). Histograms show voxelwise cross-modal correlations for each group (colored dashed lines: top 15% of negative or positive correlations for each group; black dotted lines: 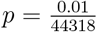, Bonferroni correction for 44318 voxels). The reported *R*^2^ indicates the proportion of variance captured by the subspace in each modality.

**Figure I.3:**
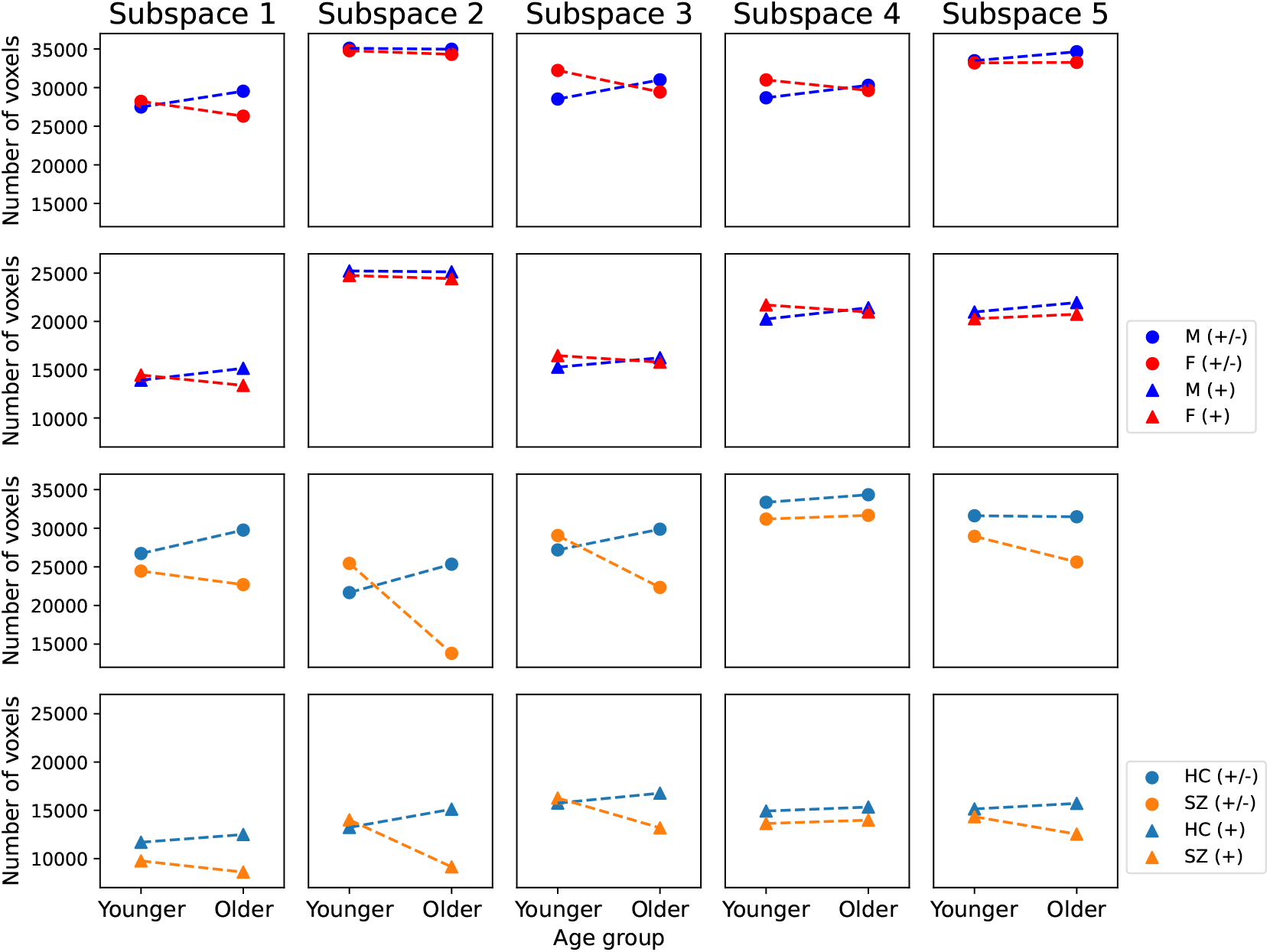
Number of voxels that show significant cross-modal correlations for age and sex groups in the UKB dataset (rows I and II), and for age and diagnosis groups in the patient dataset (rows III and IV). Rows I and III display the number of voxels with both positive and negative correlations (+*/* −), while rows II and IV display the number of voxels with only positive correlations (+). The number of voxels for older patients diagnosed with SZ is consistently less than that for their age-matched controls in four of five subspaces, implying reduced brain structure-function coupling in older patients.

## Appendix J Comparison of MSIVA *S*_2_ and MMIVA sources

### J.1 Experiments and evaluation metrics

MSIVA can be viewed as an extension of MMIVA with two main differences. First, MSIVA uses a flexible *block* diagonal subspace structure, while MMIVA uses a rigid identity matrix as the subspace structure. Second, MSIVA uses MGPCA and separate ICAs initialization, while MMIVA uses MGPCA and group ICA initialization (multimodal initialization). To further investigate similarities and differences of the recovered sources from MSIVA and MMIVA, we compared MSIVA with the subspace structure *S*_2_ and MMIVA with the subspace structure *S*_5_ through the following experiments:

1. We performed multiple linear regression (MLR) for each modality using MSIVA *S*_2_ post-CCA sources from each cross-modal subspace 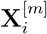 to predict each MMIVA source 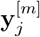:

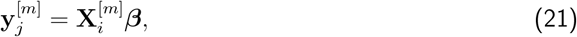

where *i* ∈ {1, …, 5} is the cross-modal subspace index in MSIVA *S*_2_, and *j* ∈ {1, …, 12} is the subspace index in MMIVA.

2. We performed multivariate analysis of variance (MANOVA) for each modality using a pair of matched MMIVA sources 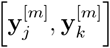 from Step 1 to predict MSIVA *S*_2_ post-CCA sources from each cross-modal subspace 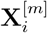:

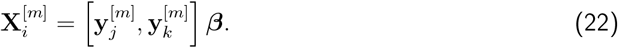

Here, (*j, k*) are a pair of matched subspace indices in MMIVA, and *i* is the cross-modal subspace index in MSIVA *S*_2_.

3. We performed brain-phenotype prediction using MSIVA *S*_2_ post-CCA sources from each crossmodal subspace, as well as each pair of matched linked 1D MMIVA sources identified in Step 1. As described in Section 2.5, we performed age prediction and sex classification tasks for the UKB dataset, as well as age prediction and binary diagnosis classification tasks (controls versus patients with SZ) for the patient dataset. We measured mean absolute error (MAE) for age regression and balanced accuracy for binary classification.

We measured the adjusted 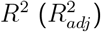 from MLR as shown below:

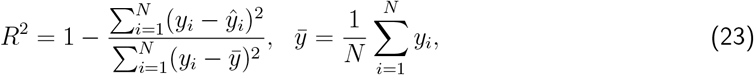

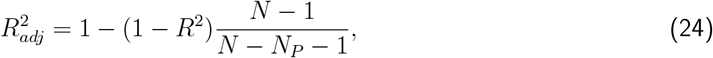

where *N* is the number of samples (here subjects) and *N*_*P*_ is the number of predictors.

### J.2 Results

Figures J.1 and J.3 show the adjusted *R*^2^ when using pairs of MSIVA *S*_2_ sources from each crossmodal subspace to predict each of the 12 MMIVA sources for the UKB dataset and the patient dataset, respectively. We reordered MMIVA sources to identify the most likely correspondence between MSIVA *S*_2_ sources and MMIVA sources. We notice that there exists some correspondence between MSIVA *S*_2_ sources and MMIVA sources. For example, for UKB sMRI data, MSIVA *S*_2_ subspace 3 sources match MMIVA source 3 (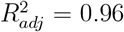), MSIVA *S*_2_ subspace 5 sources match MMIVA source 5 (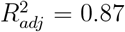), and MSIVA *S*_2_ subspace 4 sources match MMIVA source 2 (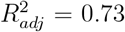). We also observe that more than two columns show high 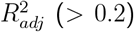 for each row, indicating that every two MSIVA *S*_2_ sources from each subspace can predict variability for more than two MMIVA sources. Note that the prediction results for fMRI are very consistent with those for sMRI.

Next, we performed MANOVA using every two matched MMIVA sources to predict the pair of MSIVA *S*_2_ sources from each cross-modal subspace. Figures J.2 and J.4 illustrate the Pillai’s trace divided by the number of modalities (*M* = 2). Note that the maximum possible diagonal Pillai’s trace value is 2 for two modalities, and dividing the Pillai’s trace by 2 shows the variance explained for each modality. We observe large off-diagonal values per column, indicating that every pair of matched MMIVA sources predicts variability of more than two MSIVA *S*_2_ sources. Note that the prediction results for fMRI are very consistent with those for sMRI.

Finally, we performed phenotype prediction using MSIVA *S*_2_ post-CCA sources from each cross-modal subspace, as well as each pair of matched MMIVA sources. Table J.1 lists the detailed prediction performance. For the UKB dataset, MSIVA sources from subspaces 5 and 4 achieved the best age regression MAE and sex classification balanced accuracy, respectively. For the patient dataset, MMIVA matched sources from subspace 2 yielded the best age regression performance, while MSIVA sources from subspaces 5 showed the best diagnosis classification performance. Overall, the linked sources from MSIVA *S*_2_ showed stronger associations with age, sex, and SZ-related effects than the paired independent sources from MMIVA.

Therefore, we conclude that MSIVA and MMIVA apportion variability to their sources in different ways. There is no perfect one-to-one mapping between MSIVA *S*_2_ sources and MMIVA sources. We also note that the mismatch appears to be more pronounced in the patient dataset than in the UKB dataset, which may be related to inherent characteristics of the patient data, such as higher population heterogeneity and smaller sample size. MSIVA linked sources from a cross-modal subspace more accurately predicted phenotype measures (age and sex in the UKB dataset and diagnosis in the patient dataset) than independent pairs of matched linked 1D MMIVA sources, indicating that higher-dimensional subspaces in MSIVA better capture phenotype-related variability than one-dimensional sources in MMIVA.

**Figure J.1:**
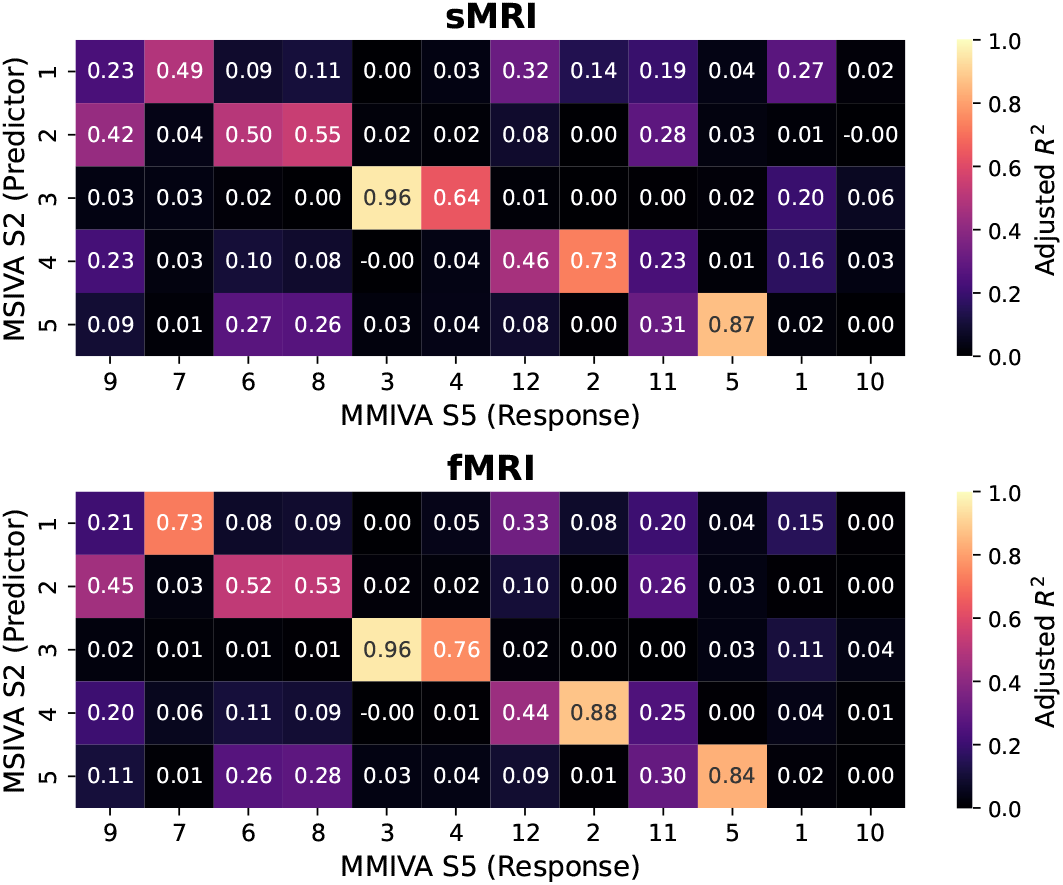
UKB neuroimaging data: Adjusted *R*^2^ using MSIVA sources to predict MMIVA sources.

**Figure J.2:**
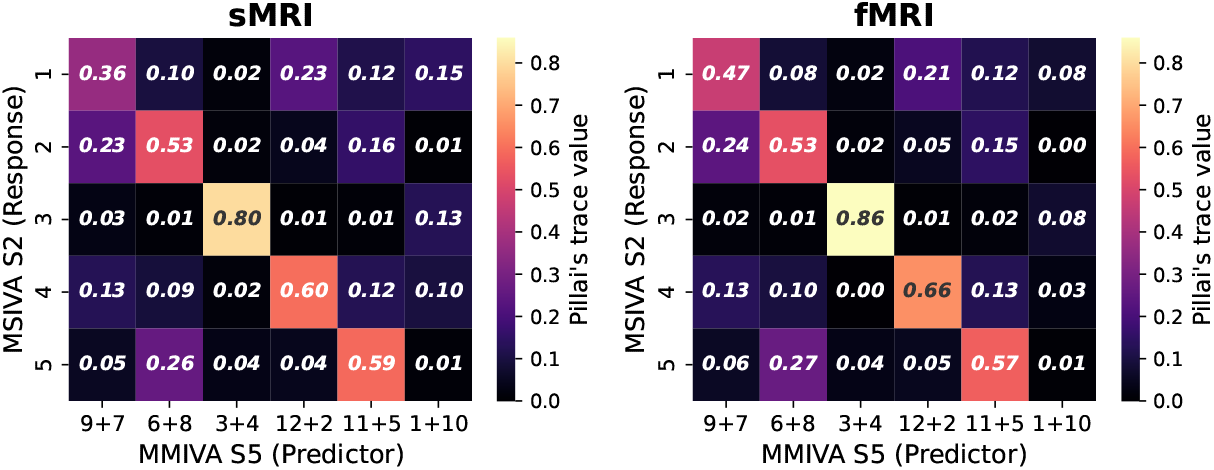
UKB neuroimaging data: Pillai’s trace value using matched MMIVA sources to predict MSIVA sources.

**Figure J.3:**
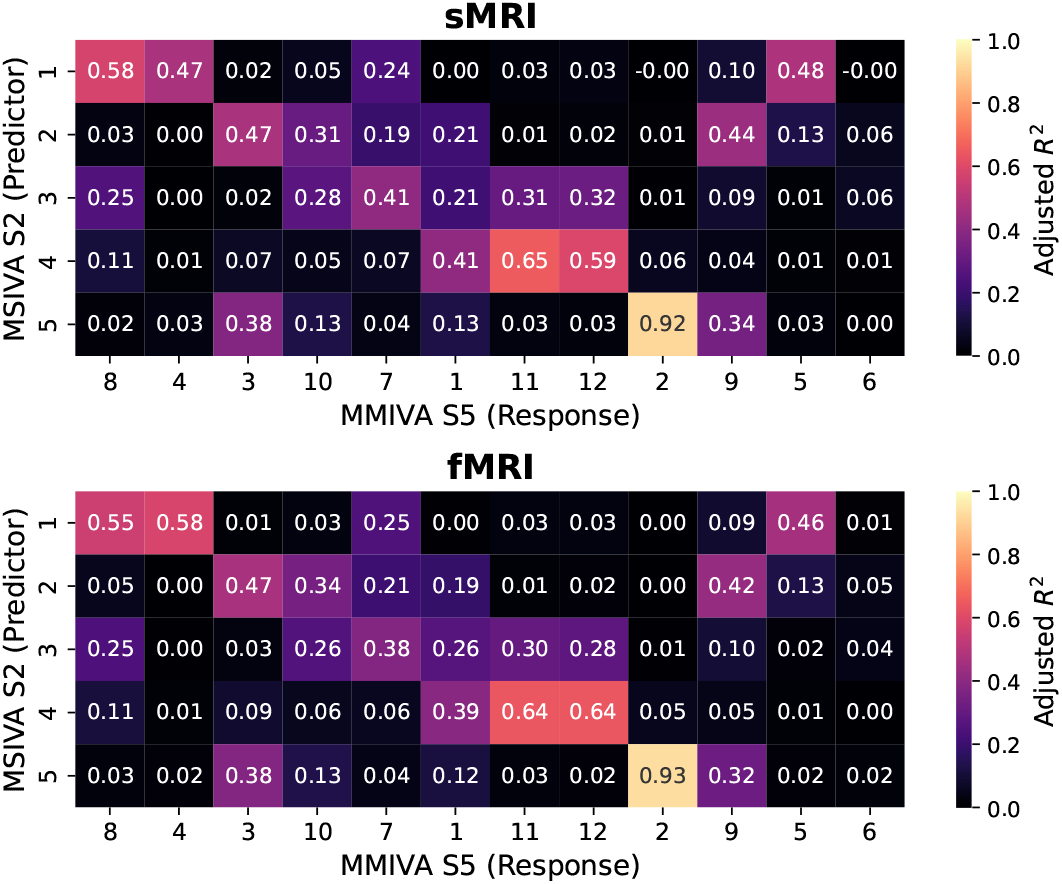
Patient neuroimaging data: Adjusted *R*^2^ using MSIVA sources to predict MMIVA sources.

**Figure J.4:**
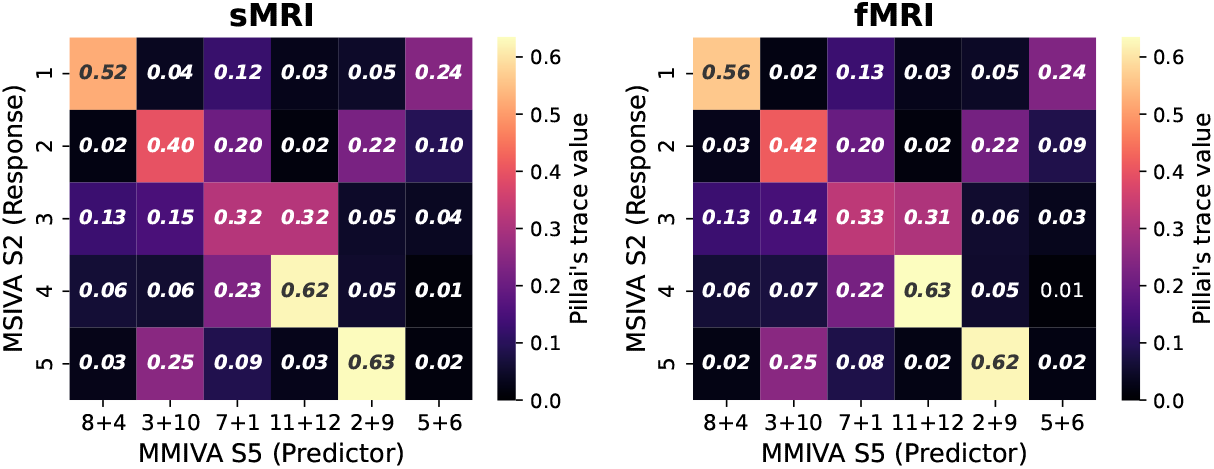
Patient neuroimaging data: Pillai’s trace value using matched MMIVA sources to predict MSIVA sources.

**Table J.1:** Phenotype prediction performance using post-CCA sources from MSIVA subspace structure *S*_2_ and matched sources from MMIVA. For the UKB dataset, MSIVA sources from subspaces 5 and 4 achieved the best age regression and sex classification performance, respectively. For the patient dataset, MMIVA matched sources from subspace 2 yielded the best age regression performance, while MSIVA sources from subspaces 5 showed the best diagnosis classification performance. Overall, the linked sources obtained by MSIVA *S*_2_ showed stronger associations with age, sex, and SZ-related effects than the paired independent sources from MMIVA. Note that we estimated sources for the UKB data and the patient data separately, thus subspaces in the UKB dataset do not correspond to those in the patient dataset.

| Subspace | 1 | 2 | 3 | 4 | 5 |
| --- | --- | --- | --- | --- | --- |
| UK Biobank dataset |  |  |  |  |  |
| MSIVA age mean absolute error (years) | 5.674 | 6.163 | 5.892 | 5.847 | **5.378** |
| MMIVA age mean absolute error (years) | 6.020 | 5.541 | 6.100 | 6.054 | 5.470 |
| MSIVA sex balanced accuracy (%) | 59.542 | 64.496 | 59.206 | **79.933** | 52.699 |
| MMIVA sex balanced accuracy (%) | 58.874 | 63.729 | 66.483 | 60.085 | 75.415 |
| Patient dataset |  |  |  |  |  |
| MSIVA age mean absolute error (years) | 10.720 | 10.470 | 11.226 | 11.445 | 10.307 |
| MMIVA age mean absolute error (years) | 10.959 | **9.822** | 10.903 | 11.679 | 10.835 |
| MSIVA diagnosis balanced accuracy (%) | 50.565 | 57.624 | 50.000 | 49.691 | **61.404** |
| MMIVA diagnosis balanced accuracy (%) | 50.752 | 57.747 | 59.919 | 50.000 | 57.695 |

## Footnotes

1 A preliminary version of this work was published at the 2023 IEEE 20th International Symposium on Biomedical Imaging (ISBI) (X. Li, Adali, et al., 2023). The present study extends the conference publication and provides a more comprehensive evaluation and analysis of the proposed method by (1) including the schizophrenia patient neuroimaging data analysis, (2) incorporating an identity matrix as an additional candidate subspace structure, (3) performing a detailed multimodal initialization comparison, (4) introducing the voxelwise brain-age delta analysis, (5) adding extensive visualizations and interpretations of the multimodal patterns and their associations, and (6) performing a quantitative comparison of loss values and other proposed metrics.

2 Stochastic gradient descent is a valid alternative approach. See Silva et al., 2020 for the derivation of the gradients.

3 We use semicolon (;) to denote that matrices are stacked vertically and comma (,) to denote that matrices are stacked horizontally.

4 The absolute values of 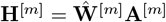 entries are reported since, in general, signs are not uniquely identifiable.

5 While a subspace is uniquely identifiable, the individual sources within each subspace are not, warranting arbitrary transformation within the subspace.

6 The modality-specific mixing matrix was estimated as the least-squares solution: 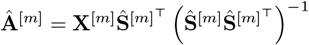.

7 For each voxel *i* and subspace *k*, we utilized singular value decomposition (SVD) across all modalities 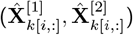 to capture the shared *reconstructed* multimodal information of each cross-modal subspace.

8 We removed mean and divided by standard deviation along the rows (subject dimension) of 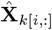.

